# Biased recruitment of H3.3 by HIRA is dictated by de-/acetylation and determines transcription memory and response

**DOI:** 10.1101/2024.08.07.606949

**Authors:** Sandra Usha Satheesan, Sourankur Chakrabarti, Amit Fulzele, Marlène Marcellin, Marie Locard-Paulet, Anne Gonzalez-de Peredo, Ullas Kolthur-Seetharam

## Abstract

Chromatin plasticity and epigenetic memory, fundamental for eukaryotic biology, are determined by differential/regulated *de novo* deposition or recycling of pre-existing histones, which in turn dictate transcriptional programs. Recruitment of the histone-H3 variant, H3.3, mediated by the HIRA chaperone complex, is both causally and consequentially associated with transcription. Despite decades of work, endogenous regulatory mechanisms that differentiate between *de novo* deposition and recycling activities of HIRA are still unknown. Here, we have investigated the pivotal role of HIRA de-/acetylation in regulating its function. Our results unequivocally establish function separation effects of acetyl and deacetyl mimic mutations of lysine-600, vis-à-vis *de novo* deposition or recycling of H3.3, respectively. Importantly, we demonstrate that HIRA deacetylation-dependent biased H3.3 recycling determines transcriptional output, possibly through preferential enrichment of H3.3-K36me3. Besides unraveling tunable regulatory mechanism that governs HIRA function, we illustrate a causal link between the chaperone activity, biased recruitment of pre-existing histones, and gene expression.

## Introduction

Histone variants and post-translational modifications (PTMs) that form the nucleosome, which is the structural and functional unit of chromatin, have been well-established as being pivotal for all DNA transactions within the nucleus [1–4]. Notably, the repertoire of compositional and functional diversity of chromatin generated by histone variants and their PTMs has a deterministic impact on the nucleosome and thus all of eukaryotic biology [2, 3, 5–7]. Moreover, chromatin plasticity and transcriptional memory, dependent on nucleosomal dynamics, are crucial for the establishment and maintenance of cellular and functional identities besides regulating inducible gene expression [8, 9].

Histone H3 and its variants are known to play a deterministic role in transcription, replication, and repair, in addition to dictating heterochromatin and euchromatin dynamics [10–12]. H3.1 and H3.2, the canonical H3 variants, are synthesized and incorporated into the chromatin in a replication-dependent manner [11]. Transcription-dependent replacement of H3.1/3.2 by H3.3 is critical for gene expression regulation [13–16]. This replication-independent deposition of H3.3 is locus-specific and therefore coupled to the rate/extent of transcription of specific genes [17, 18]. Despite this, our understanding of regulatory mechanisms that control differential recruitment of H3.3, which is linked to divergent transcriptional output across loci, is still poor.

Seminal studies identified the HIRA complex, with HIRA, UBN1, and CABIN1 as core components of the machinery that recruits/deposits H3.3 onto chromatin [19–21]. Deletion analysis of each of these factors leads to a complete loss of activity of the entire complex largely due to destabilization and degradation of the proteins themselves [21–23]. While this has clearly demonstrated their interdependence, the exact molecular functions of HIRA, UBN1, and CABIN1 within the complex have remained elusive. A recent report that employed mutant versions of HIRA, which affects interaction with UBN1 and ASF1a, has shown that it is required for depositing H3.3 [24]. Nonetheless, the debate over the role of HIRA and the other interacting proteins in orchestrating the chaperone activity to regulate spatiotemporal deposition of H3.3 is still largely unresolved.

Recruitment of H3.3 at actively transcribing loci could be brought about either by depositing freshly synthesized H3.3 (*de novo*) or recycling pre-existing H3.3. It is important to note that the HIRA complex is required for both *de novo* deposition and recycling of H3.3 [24]. A recent report showed that while HIRA-ASF1 complexation mediates the recycling of H3.3, HIRA trimerization and its interaction with UBN1 is required for *de novo* deposition [24]. However, mechanisms that differentiate these two activities of the HIRA protein itself or govern the dynamics of functionally distinct complexes are still unknown. Specifically, regions/sites within HIRA and regulatory modifications that allow function separation, which is integral to initiating and sustaining gene expression need to be uncovered.

Among others, reversible post-translational modifications (PTMs) are well known to regulate protein structure, interactions, and activities/functions [2, 5, 25–27]. In this regard, very little is known about the PTM-dependent regulation of HIRA, except for phosphorylation and glycosylation, which were shown to affect cellular differentiation and senescence [28–31]. Although global proteomics-based approaches have identified acetylation and SUMOylation of HIRA [32–34] if/how they impinge on HIRA functions needs to be investigated. Moreover, it is important to uncover whether any of these modifications exert regulatory control over *de novo* deposition and recycling activities of HIRA and the consequent impact on gene transcription.

In the present study, we identify an acetylation-dependent functional bias in HIRA activity of *de novo* H3.3 deposition and recycling. By employing acetylation (K-Q) and de-acetylation (K-R) mimics of HIRA we demonstrate that the acetyl mimic has enhanced *de novo* deposition of H3.3 while the deacetyl mimic has enhanced recycling and the differential activities. Furthermore, to elucidate the downstream consequences, we employed various transcriptional paradigms to dissect how these differential HIRA activities impact gene expression.

## Results

### Mammalian HIRA is acetylated

As described earlier, little is known about HIRA complex regulation that is brought about by post-translational modifications. Among the PTMs, acetylation is especially interesting because it is responsive to cellular metabolic status [35, 36]. Consistent with earlier reports that identified HIRA acetylation [32, 34], our efforts at unraveling SIRT1- and fed-fast dependent global acetylproteome also revealed HIRA to be acetylated in mice (Fig. 1A). Importantly, the lysine-599, which we found to be acetylated in mice, is conserved across vertebrates (Fig. 1B). Notably, acetylation at this site has been reported by an earlier study, which characterized SIRT1-dependent acetylproteome in mouse embryonic fibroblast (MEF) [32]. Together, this study and our acetyl proteome data clearly highlight SIRT1-dependent HIRA acetylation.

**Fig. 1.**
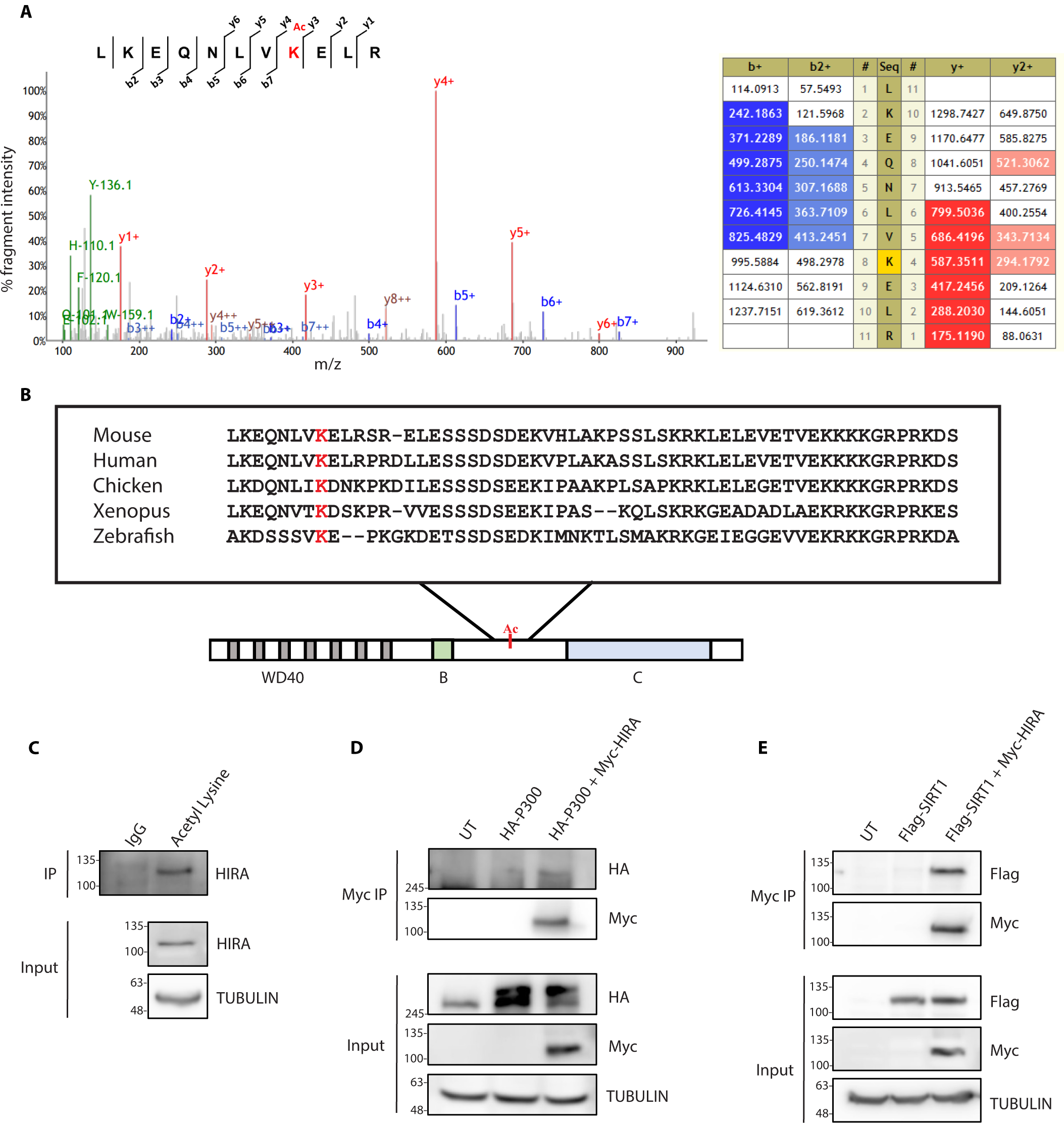
Mammalian HIRA is acetylated. **(A)** Representative fragmentation mass spectrum of HIRA peptide 592-602 (protein sequence Q61666-1 in Uniprot), obtained from a precursor peptide of mass 1410.819 Da, showing an increment of 42Da corresponding to an acetylation. Red peaks correspond to the y serie of N-terminal fragments and blue peaks to the b serie of C-terminal fragments obtained by high-energy collision induced dissociation, as illustrated on the inserted peptide sequence. The table on the right shows the masses of the different fragment ions detected from the mass spectrum by the search engine software Mascot. Fragments y4, y5 and y6 are indicative of an acetylation site on the lysine at position 8 of the peptide sequence, corresponding to lysine-599. **(B)** Schematic showing HIRA acetylation and protein sequence alignment depicting K-599 conservation in vertebrates. **(C)** Immunoprecipitation (IP) assay performed using pan-acetyl lysine antibody followed by western blot using HIRA antibody to detect acetylated HIRA in mouse liver (N=2). **(D)** HEK293T cells co-transfected with HA-P300 and Myc-HIRA followed by coimmunoprecipitation (coIP) assay using Myc beads and western blot analysis performed using HA antibody to identify the interaction between HIRA and P300 (N=2). **(E)** HEK293T cells co-transfected with Flag-SIRT1 and Myc-HIRA followed by coIP assay using Myc beads and western blot analysis performed using Flag antibody to identify interaction between HIRA and SIRT1 (N=2).

Next, we wanted to biochemically validate HIRA acetylation, and towards this, we performed an immunoprecipitation assay using pan-acetyl lysine antibody, and western blotting using HIRA antibody and found HIRA to be acetylated in mouse liver (Fig. 1C). Further, as expected, human HIRA protein in HEK293T cells was also found to be acetylated (fig. S1A).

Protein acetylation is reversibly regulated by lysine acetyltransferases (KATs) and lysine deacetylases (KDACs) [27]. While data from the literature indicate p300/CBP as KATs that potentially acetylate HIRA [34], we wanted to check if other KATs are also involved in HIRA acetylation. As shown in fig. S1B, overexpression of P300 but not PCAF and GCN5 led to an increase in HIRA acetylation. Corroborating the involvement of p300 in acetylating HIRA, immunoprecipitation assays clearly showed that these two proteins interact with each other in HEK293T cells (Fig. 1D). Given that HIRA acetylation was found to be sensitive to the presence or absence of SIRT1, we wanted to check if they formed a complex. Co-immunoprecipitation assays clearly showed that SIRT1 and HIRA interacted with each other (Fig. 1E), and substantiated the results described above.

### Acetylation alters HIRA chromatin association

Next, we wanted to study the functional consequence, if any, of HIRA acetylation at lysine-600 (K600). Lysine to glutamine (K-Q) and lysine to arginine (K-R) mutations are well-established in the literature to mimic acetylated and deacetylated proteins, respectively [37–40], due to the similarity in charge and chemical structure. Hence, we cloned K600Q and K600R mutants of human HIRA as described. Upon transient transfection in HEK293T cells, WT and the mutants showed similar expression levels suggesting that the mutation did not affect stability (fig. S2A). However, we wanted to ascertain if acetyl and deacetyl mimics of HIRA impinged on localization and chromatin association. Ectopically expressed WT and mutant HIRA proteins were probed for in whole-cell lysate, cytoplasm, nucleoplasm, and chromatin fractions (Fig. 2A and fig. S2B). Even though we observed the ectopically expressed HIRA proteins in the cytoplasm (fig. S2B), possibly an artifact of overexpression, they were abundantly found in the nuclear fractions (Fig. 2A), as expected. Comparing levels of WT and mutant HIRA in nucleoplasmic and chromatin fractions clearly indicated that the charge state of K600 affected chromatin association (Fig. 2A).

**Fig. 2.**
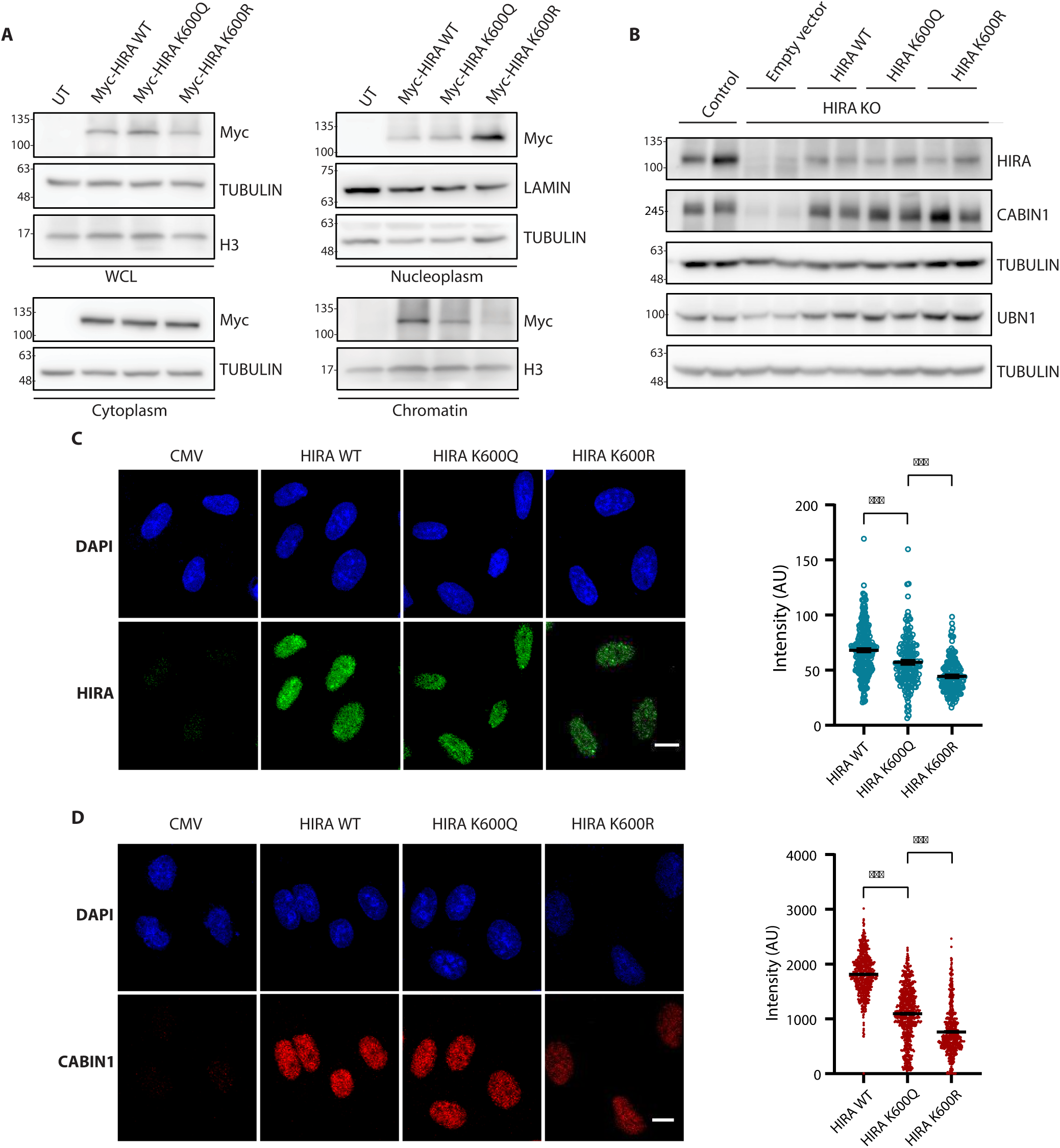
Acetylation alters HIRA chromatin association. **(A)** HEK293T cells transfected with Myc-HIRA WT, K600Q, and K600R followed by biochemical fractionation obtaining the fractions WCL, cytoplasm, nucleoplasm, and chromatin. Myc-HIRA levels and markers of respective fractions assayed using western blot analysis (N=3). **(B)** Representative western blots for levels of HIRA, CABIN1, and UBN1 in control, HIRA KO, and HIRA KO cells rescued with HIRA WT, K600Q, and K600R (n=2, N=2). **(C-D)** Representative images for chromatin-bound HIRA **(C)** and CABIN1 **(D)** levels in HIRA KO cells rescued with HIRA WT, K600Q, and K600R using immunofluorescence assay and the quantification (right). The data presented is mean ± S.E.M. (N=2, n>150) with the individual points shown. Statistical significance was calculated using one-way ANOVA with Tukey’s test for multiple comparisons between groups (*, p < 0.05; **, p < 0.01; ***, p < 0.001). Scale bars −10 µm.

Notably, while there was no difference between WT and K600Q mutant, K600R HIRA was more abundant in the nucleoplasm and not on the chromatin (Fig. 2A). To probe this further, we subjected the chromatin fraction to differential salt extractions (fig. S2C). As shown in fig. S2C chromatin-bound K600R HIRA was dramatically less under high salt conditions corroborating lower affinity to chromatin. Notwithstanding the artifacts associated with overexpression, these results indicated that de-/acetylation led to differential binding of HIRA to the chromatin.

To provide a physiologically relevant assessment of HIRA de-/acetylation, in addition to negating artifacts of overexpression, we utilized HIRA knock-out cells that were stably transfected with WT, K600Q, and K600R HIRA to restore its expression (Fig. 2B and fig. S2D). Specifically, Hela-HIRA CRISPR knock-out cells [23] were used and we found that these stable lines had comparable levels of expression of HIRA (Fig. 2B). As reported earlier, loss of HIRA resulted in depletion of CABIN1 and UBN1 proteins which were restored in WT HIRA rescue cells (Fig. 2B). However, it is important to note that CABIN1 and UBN1 expression were similarly restored in cells stably expressing K600Q and K600R mutants of HIRA (Fig. 2B). Besides validating the rescue, this also suggested that de-/acetylation of HIRA did not perturb the stabilities of CABIN1 and UBN1. Having established the rescue cell lines that expressed WT and mutant forms of HIRA to endogenous levels, we wanted to confirm if deacetylation indeed leads to lower chromatin affinity. As detailed in the methods section, Triton X-100 extraction followed by fixation of cells, to deplete soluble and/or loosely chromatin-bound proteins, clearly showed lower levels of K600R HIRA that remained associated with chromatin (Fig. 2C). Furthermore, we probed for CABIN1 localization and found similar results (Fig. 2D). In addition to corroborating the results described above this illustrates the effect of deacetylation vis-à-vis chromatin association of HIRA complex.

### HIRA complexation is independent of acetylation

Interaction of HIRA with CABIN1 and UBN1 is essential for the functional complex and an absence of HIRA is well established to destabilize the complex and induce degradation of CABIN1 and UBN1 [21–23]. Given this, restoration of CABIN1 and UBN1 expression in Hela rescue cells (WT, K600Q, and K600R) (Fig. 2B) indicated that de-/acetylation mimic mutants do not affect complexation. However, to provide confirmatory evidence we assayed for interactions of HIRA with CABIN1, UBN1, and ASF1A under endogenous and overexpression conditions as depicted (Fig. 3A-C). ASF1A, although not necessary for HIRA-dependent deposition of H3.3 at transcriptionally active loci, is known to interact with HIRA and has been speculated to be necessary for binding of H3.3 dimers to the HIRA complex [41–43].

**Fig. 3.**
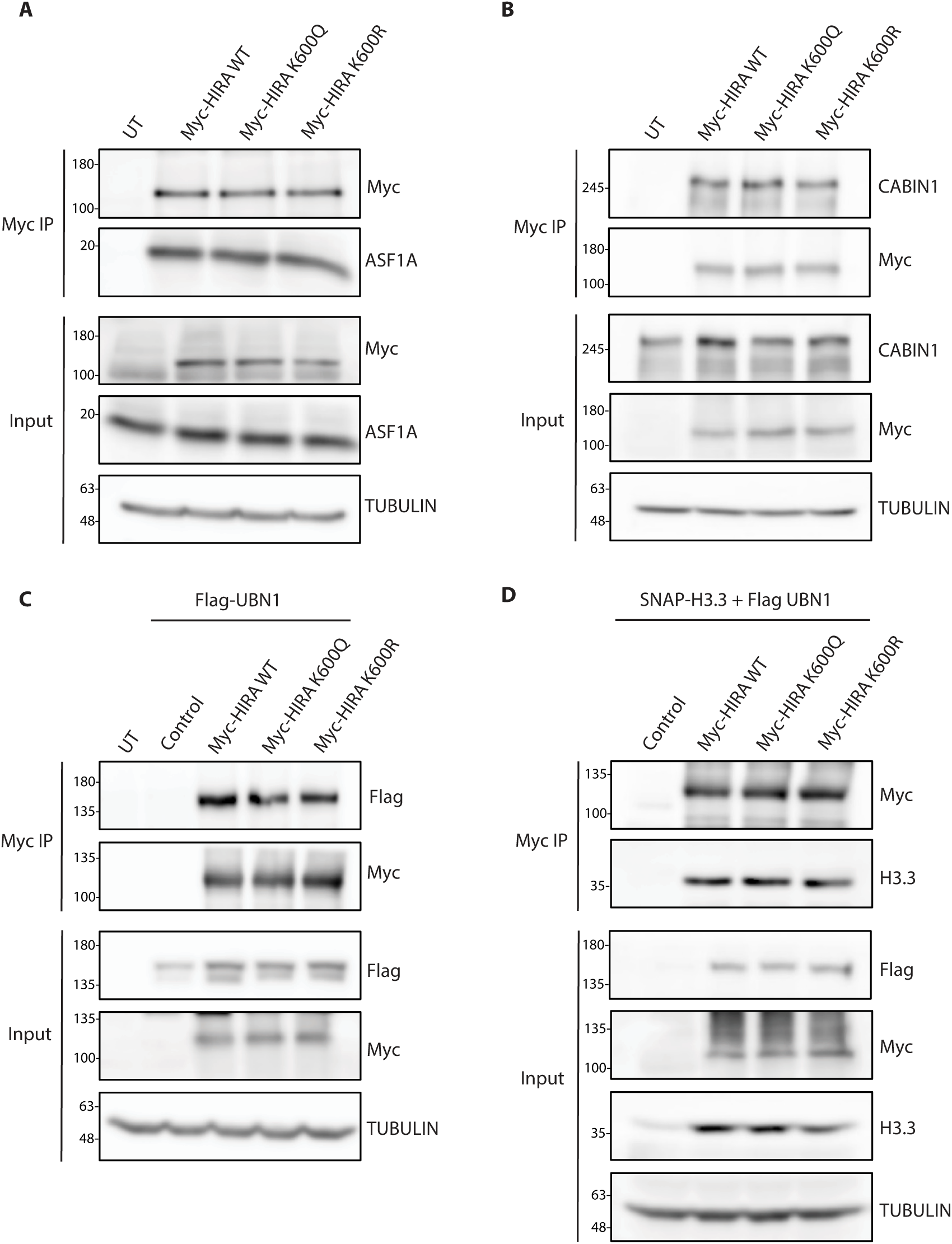
HIRA complexation is independent of acetylation. **(A-B)** HEK293T cells transfected with Myc-HIRA WT, K600Q, and K600R followed by coIP assay using Myc beads and western blot to assay for interactions with **(A)** ASF1A and **(B)** CABIN1 (N=3). **(C)** HEK293T cells co-transfected with Flag-UBN1 and Myc-HIRA WT, K600Q, and K600R followed by coIP assay using Myc beads and western blot for Flag to assay for interaction with Flag-UBN1 (N=2). **(D)** HEK293T cells co-transfected with Flag-UBN1, SNAP-H3.3, and Myc-HIRA WT, K600Q, and K600R followed by coIP assay using Myc beads and western blot for H3.3 to assay for interaction with SNAP-H3.3 (N=3).

We carried out a coimmunoprecipitation assay using Myc-tagged WT, K600Q, and K600R HIRA proteins, from HEK293T cells and found no difference in their interaction with endogenous ASF1A (Fig. 3A) and CABIN1 (Fig. 3B). Since our attempts to detect interaction with endogenous UBN1 were futile, we made use of the FLAG-UBN1 expression vector and found that the interactions of ectopically expressed UBN1 with Myc-HIRA WT, K600Q, and K600R remain unaltered (Fig. 3C). A recent study, which used HIRA mutations within domains that are either necessary for trimerization or its interaction with UBN1 and ASF1, characterized sub-molecular complexes with exclusive recycling or *de novo* activities [24]. Importantly, unlike this, we found de-/acetylation of HIRA, which is physiological, did not affect complexation with ASF1A, CABIN1, and UBN1. Reverse immunoprecipitation wherein ASF1A and UBN1 were used as baits to pull down the complex confirmed these results (fig. S3A and B). Further, to check for the strength of interactions, immunoprecipitations were done under high salt conditions as indicated, which showed no change in binding affinities (fig. S3C-F). In addition to this, we also wanted to ascertain if the de-/acetylation status of HIRA impacted binding with H3.3. Since an earlier report illustrated that overexpressing UBN1 was necessary for capturing the interaction between HIRA and H3.3 [22] we used the same strategy to evaluate the impact of de-/acetyl mimic mutants. As shown in Fig. 3D we found no change in the interaction of H3.3 with the HIRA complex. Together, these clearly suggested that the acetylation status of HIRA does not perturb its ability to bind/interact with its partners, which are critical for chaperone activity.

### Acetylation dependent bias in HIRA activity of *de novo* H3.3 deposition and recycling

Despite extensive characterization of the HIRA complex, active degradation of the core components in the absence of HIRA has confounded findings, which ascribe specific activities or roles to each of these proteins [21–23]. Nonetheless, the complex itself has been unequivocally established to be essential for both *de novo* deposition and recycling of H3.3 [24, 41, 44, 45]. In the absence of any evidence vis-à-vis regulatory mechanisms that either bias or switch between *de novo* and recycling activities of the HIRA complex, we were prompted to investigate the importance of de-/acetylation of the HIRA protein in this regard.

The SNAP-tag approach has been employed as a method of choice to investigate histone dynamics, both recruitment and turnover [46]. In particular, kinetic paradigms of pulse/chase (detailed in the methods section) have allowed selective visualization and quantification of either newly synthesized/deposited or recycled histones. Therefore, we cloned human H3.3 as a fusion construct with a SNAP-tag, at its C-terminus (H3.3-SNAP). Following validation of this clone and its expression, we stably transfected HIRA CRISPR KO HeLa cells to express H3.3-SNAP (fig. S4A). Despite our attempts, we failed to generate cells that have stable expression of both H3.3-SNAP and HIRA protein. Given this, HIRA KO HeLa cells stably expressing H3.3-SNAP were transfected with WT, K600Q, and K600R constructs to restore HIRA (fig. S4B and C), which we have earlier shown to reinstate HIRA/CABIN1/UBN1 complex (Fig. 2B). This also ensured that there was no variability vis-à-vis the abundance of H3.3-SNAP and allowed us to unbiasedly compare the *de novo* deposition and recycling activity of the HIRA complex, as a consequence of de-/acetylation.

As described earlier and in the methods section, cells were treated with SNAP-Cell Block reagent to quench pre-existing H3.3-SNAP and subsequently pulsed with SNAP-Cell TMR-Star to specifically label newly synthesized/deposited H3.3-SNAP, as indicated (Fig. 4A and fig. S4B). On quantifying TMR fluorescence, we found *de novo* deposition of H3.3-SNAP in cells that expressed WT, K600Q, and K600R mutants, which was anticipated (Fig. 4B). However, it is important to note that there was a significant increase in *de novo* deposition in cells that expressed K600Q HIRA, over and above the WT and K600R mutant, which were otherwise comparable (Fig. 4B).

**Fig. 4.**
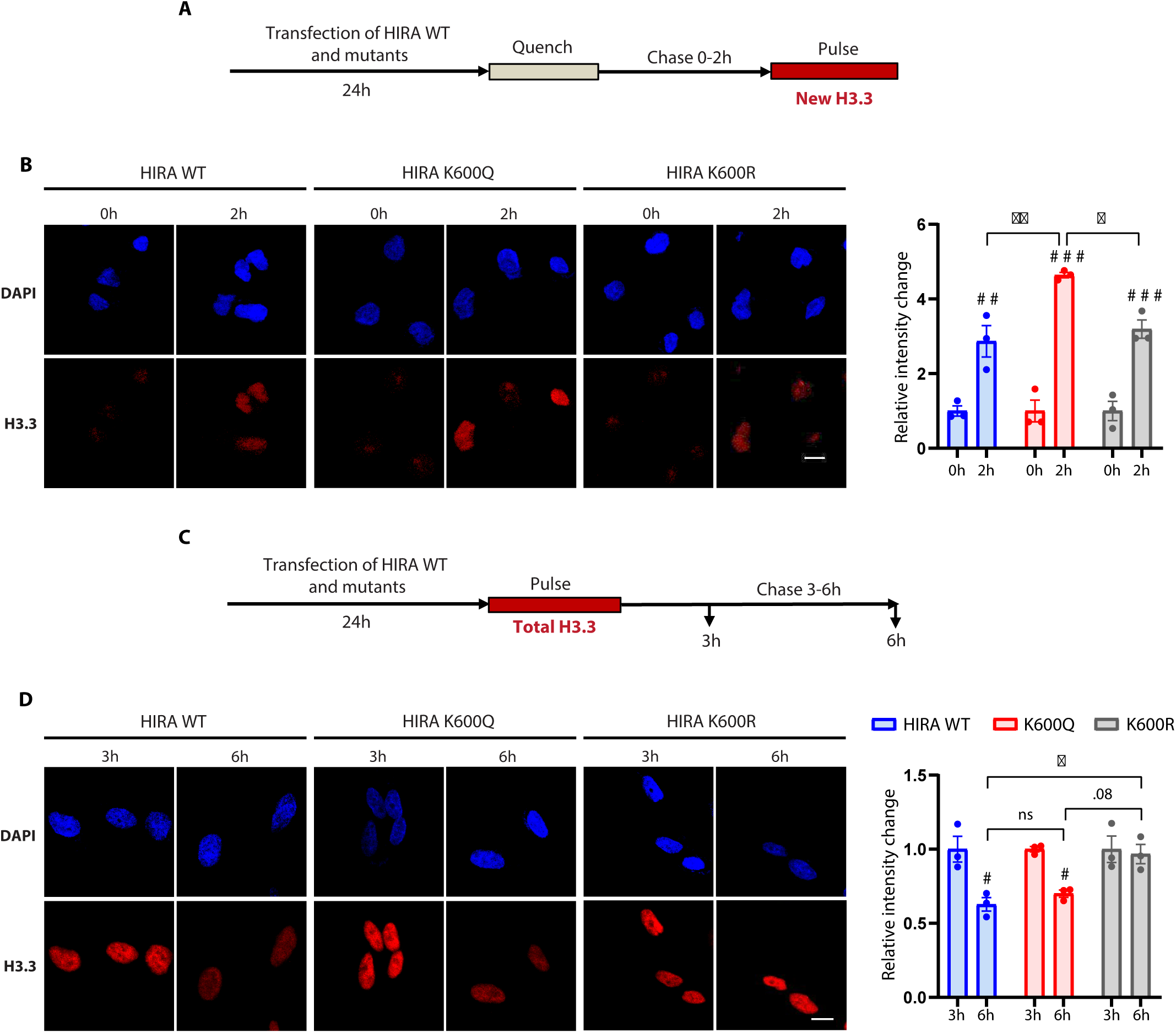
Chaperone activity of HIRA is governed by de-/acetylation. **(A)** Schematic depicting the experimental paradigm for assessing H3.3 *de novo* deposition. **(B)** Representative immunofluorescence images showing H3.3-SNAP (red) and DAPI (blue) in Hela-HIRA-KO cells stably expressing SNAP-H3.3. Cells were transfected with Myc-HIRA WT, K600Q, and K600R followed by *in vivo* quench-chase-pulse labeling of newly synthesized and deposited H3.3-SNAP during the 2h chase window. **(C)** Schematic depicting the experimental paradigm used for assessing H3.3 recycling. **(D)** Representative immunofluorescence images showing H3.3-SNAP (red) and DAPI (blue) in Hela-HIRA-KO cells stably expressing SNAP-H3.3. Cells were transfected with Myc-HIRA WT, K600Q, and K600R followed by *in vivo* pulse labeling of pre-existing H3.3-SNAP and a chase period of 3h and 6h. The quantifications represented as mean ± S.E.M. (N=3, n=3 (150-200 cells counted per coverslip)). Statistical significance was calculated using two-way ANOVA with Tukey’s test for multiple comparisons between groups. #, p < 0.05; # #, p < 0.01; # # #, p < 0.001 indicate statistical significance across different time points within HIRA WT, K600Q, and K600R. *, p < 0.05; **, p < 0.01; ***, p < 0.001 indicate statistical significance for the same time points between HIRA WT, K600Q, and K600R. Scale bars-10µm.

Next, to check the impact on recycling activity, cells were pulsed with TMR to label pre-existing H3.3-SNAP and chased to score for retention/loss of signal as an indicator of recycling (Fig. 4C and fig. S4C). To further negate experimental variabilities, we used the signal at the 3h timepoint in each of these respective cell lines, as a baseline and quantified the reduction over a 3h window (Fig. 4D). It was interesting to note that while there was a substantial reduction in H3.3-SNAP labeled with TMR in cells that expressed WT and K600Q mutant, there was no change in K600R HIRA expressing cells (Fig. 4D). To further substantiate this, we over-expressed WT and mutant HIRA in normal HIRA cells and scored for recycling activity as described (fig. S4D). We found that K600R HIRA displayed better recycling activity when compared to WT and K600Q mutant (fig. S4E). Together, these results clearly suggested that acetyl and deacetyl mimics of HIRA have biased *de novo* deposition and recycling activities.

### HIRA acetylation impinges on gene expression

Even though, HIRA’s role in H3.3 recruitment/deposition is well established, causal evidence linking *de novo* deposition and recycling activities to transcription is still lacking. Based on our result described above it was obvious for us to investigate this link using de-/acetylation mutants of HIRA. Towards this, we used multiple paradigms to induce gene expression in response to heat shock and IFN-γ treatments in cells that were either stably or transiently transfected with WT, K600Q, and K600R mutants, as indicated (Fig. 5A-F and fig. S5A). Consistent with earlier reports [47, 48] exposure to 42°C led to a robust induction of transcription of key heat shock proteins (Fig. 5B and fig. S5A). However, we found a significant difference between the WT and the mutants especially in the case of K600R, which displayed maximal expression of *Hsp70* and *Hsp27* (Fig. 5B).

**Fig. 5.**
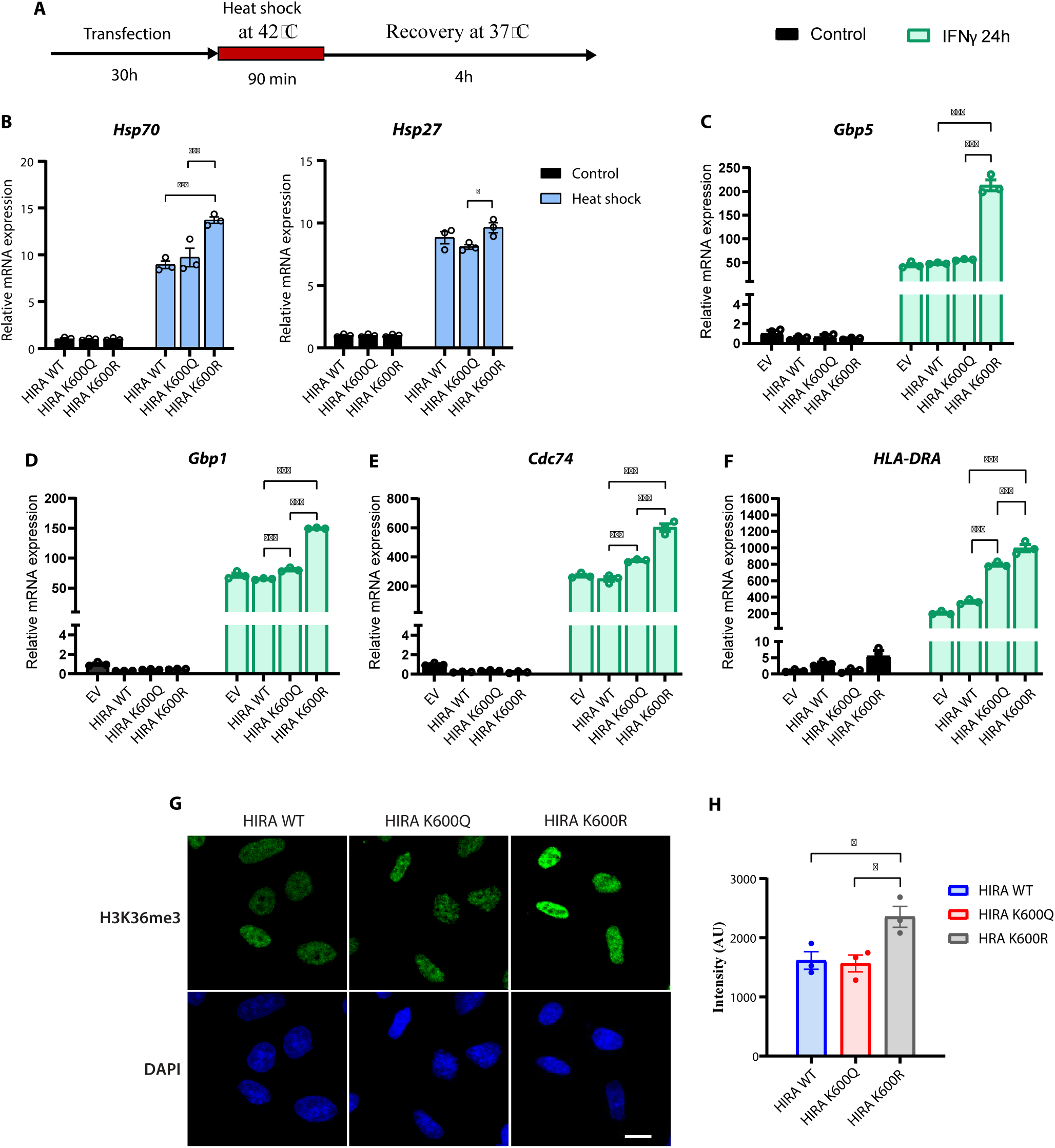
Deacetyl mimic HIRA impinges on extent of gene expression and activatory epigenetic marks. **(A)** Schematic depicting the heat shock experimental paradigm. **(B)** Relative expression of *Hsp70* and *Hsp27* mRNAs in HEK293T cells transfected with HIRA WT, K600Q, and K600R followed by the heat shock paradigm. mRNA levels were normalized to actin (*Actb*) and plotted as fold change with respect to the respective control conditions (no heat shock). **(C-F)** Relative expression of *Gbp5* **(C)**, *Gbp1* **(D)**, *Cdc74* **(E)**, and *HLA-DRA* **(F)** mRNAs in HIRA KO cells stably expressing empty vector (EV), HIRA-WT and K600Q and K600R treated with IFNγ for 24 hrs. mRNA levels were normalized to actin (*Actb*) and plotted as fold change with respect to the control condition (no IFNγ treatment) of EV. **(G-H)** Representative images for chromatin-bound H3K36me3 levels in HIRA KO cells rescued with HIRA WT, K600Q, and K600R using **(G)** immunofluorescence assay and **(H)** its quantification. Data represented as mean ± S.E.M. (N=2, n=3). Statistical significance was calculated using two-way ANOVA with Tukey’s test for multiple comparisons between groups (*, p < 0.05; **, p < 0.01; ***, p < 0.001). Scale bars −10 µm.

Next, to negate artifacts of overexpression we utilized HIRA KO cells that were stably rescued with WT, K600Q, and K600R mutants. Earlier studies have shown IFN-γ treatment of HeLa cells to result in active transcriptional upregulation of the downstream genes [49], which was recapitulated in our cells (Fig. 5C-F). It was indeed intriguing to find that rescuing HIRA CRISPR KO cells with HIRA WT did not result in enhanced expression of IFN-γ target genes. We surmised that this is possibly due to either a compensatory mechanism or because of the steady state effect that potentially masks kinetic differences in transcription. Despite this caveat, it is interesting to note that cells that were rescued with K600R showed heightened expression of *Gbp1*, *GBP5*, *HLA-DRA*, and *Cdc74*, more than what was observed in the case of WT and K600Q (Fig. 5C-F). Although not surprising, these responses were not associated with bulk changes in global transcription or phosphorylated form of RNA Pol-II (fig. S5B-G). Hence, taking together the heat shock and IFN-γ induced gene expression, our results demonstrate that the deacetylated form of HIRA is necessary for heightened induction of transcription.

Since acetylation and deacetylation mimic mutants elicited differential effects with regards to both H3.3 recruitment/deposition and transcription, we wanted to check if this was associated with distinct patterns of epigenetic modifications. This is important since newly synthesized H3.3 and recycled H3.3 will expectedly have differential post-translational modifications [50, 51]. It was interesting to note that H3.3 associated H3K36me3, which is a diagnostic marker of rate/extent of transcription [52–55], showed a robust increase in HIRA CRISPR KO cells that were rescued with K600R HIRA (Fig. 5G and H). Besides complimenting the data on transcriptional change, this illustrated that deacetylated HIRA plays a pivotal role in coupling or facilitating transcription via H3.3 recycling.

## Discussion

Chromatin remodeling and nucleosomal composition are pivotal for gene expression regulation and often rate-limiting [56–58]. Independent of mechanisms involved in nucleosomal disassembly and reassembly, factors that determine the deposition of histone variants are well known to dictate transcriptional output globally and, in a gene-specific manner [11, 57, 59–66]. Moreover, the recruitment of newly synthesized or recycled pre-existing variants, along with epigenetic marks, is a key component of transcriptional memory and plasticity [67–69]. Even though histone chaperones are known to mediate histone variant recruitment, specifically of H3.3 by HIRA [15, 44, 61], regulatory mechanisms that not only distinguish *de novo* deposition versus recycling but also couple this to transcription have remained elusive. In this regard, our study has uncovered how HIRA acetylation and deacetylation act as a function separation modulatory mechanism vis-à-vis H3.3 deposition, which is essential for transcription and hitherto unknown.

Reversible post-translational modifications (PTMs) provide precise mechanistic control over protein structure and function, including via complexation, localization, and turnover [2, 5, 25–27]. This is even more apparent in nuclear and chromatin factors that orchestrate gene expression [2, 5, 25–27, 70]. Given the abundance of lysine acetyltransferases and deacetylases in the nucleus that impinges on histones, transcription factors, and chromatin remodelers, it is intuitive to hypothesize histone chaperones to be similarly regulated. Even though HIRA has been identified in global acetyl proteome analysis the importance of HIRA acetylation has not been investigated thus far [32, 34]. Consistent with earlier reports [32, 34] we identified HIRA acetylation at lysine 599, which is conserved across species. In addition, our results also demonstrate that acetylation and deacetylation dynamics are regulated by CBP/P300 and SIRT1, respectively. This is not surprising because these factors are known to impinge on both locus-specific and global de-/acetylation-dependent transcription [71–74].

As mentioned earlier, interactions between HIRA, CABIN1, and UBN1 leading to the formation of a trimeric complex are essential for not only the stability of these proteins [21–23] but also to bring about H3.3 deposition [24, 41, 44, 45]. It is important to note the molecular functional identities of each of these components have not been deciphered until now except for a very recent study that utilized point mutants of HIRA that abolish its interactions with ASF1 and UBN1. Specifically, findings from this study ascribed domain dependence, within HIRA, for mediating H3.3 recycling or *de novo* deposition [24]. Since these mutants of HIRA are not expressed endogenously, mechanisms that differentiate molecular functions of HIRA with respect to H3.3 recycling and *de novo* deposition remain largely unknown. Given this, we have discovered HIRA acetylation as a regulatory modification without affecting its ability to complex with UBN1, CABIN1, and ASF1A. Using classical approach to endogenously express acetyl and deacetyl mimics of HIRA we clearly illustrate function separation effects mediated through lysine 600. Our results establish that while acetylation favors *de novo* deposition of H3.3, deacetylation is necessary for recycling of H3.3, which was hitherto unknown.

Even though it is evident that the modifications on newly synthesized and previously deposited H3.3 are different [50, 51], if or how these epigenetics marks are differentially employed for *de novo* deposition and recycling remains to be addressed. Conversely, it is anticipated that histone modifications that are associated with active transcription would be enriched in the case of biased recycling activity of the HIRA complex. Consistent with this hypothesis we have indeed found a robust global increase in H3K36me3 in rescue cells that express the deacetyl mimic of HIRA. This provides confirmatory evidence vis-à-vis enhanced recycling activity of deacetylated HIRA, which was unknown until now. This is important since the function separation deacetyl mimic of HIRA will enable future efforts aimed at elucidating histone marks that are distinctly employed by HIRA for recycling H3.3.

Even though our study unequivocally establishes acetylation dependent functional bias in HIRA activity, histone PTMs that constitute this molecular distinction of pre-existing and newly synthesized H3.3 by de/-acetylated HIRA needs to be unraveled in the future. Another intriguing observation is that HIRA acetylation governs chromatin association of the complex, but if/how this leads to differential *de novo* and recycling activity, and consequent transcriptional changes remains unknown. In the absence of studies that have addressed transience of chromatin bound HIRA, it is tantalizing to propose de-/acetylation dependent HIRA residence on actively transcribing loci that contributes to differential recognition of pre-existing H3.3.

Albeit the importance of H3.3 deposition as a key determinant of transcriptional output is well known [15, 17, 18, 75, 76], it is surprising to note that there is a paucity of information with regards to the recruitment of freshly synthesized or recycled H3.3. While it is intuitive to expect that recycling H3.3 would lead to enhanced transcription this has not been evidenced thus far. This is critical because the ratio metric abundance of new versus recycled H3.3 forms the basis for transcriptional plasticity and memory. In this context, we establish that biased recycling is associated with a graded increase in the transcription of inducible genes. Moreover, when coupled with HIRA acetylation status we demonstrate a regulatory mechanism that tunes gene expression through differential recruitment of H3.3.

## Materials and Methods

### Plasmids and constructs

Human H3.3 open reading frame (ORF) was amplified by PCR (from the cDNA prepared from HEK293 cells), and cloned between the AscI and EcoRI sites of the pSNAP_f_ vector (NEB) to generate H3.3-SNAP plasmid. Human HIRA ORF (3054 nucleotides) was amplified by PCR (from the cDNA prepared from HEK293 cells), and cloned between the EcoRI and SalI sites of the pCMV-Tag 3B vector to generate pCMV-Tag 3B HIRA WT plasmid. Lysine 600 to glutamine and arginine, K600Q and K600R mutants respectively, were made using site directed mutagenesis wherein primers with specific point mutations were used in PCR to amplify mutant 5’ fragment. The 5’ mutant fragments were cloned between EcoRI and BbvCI sites of pCMV-Tag 3B HIRA WT plasmid following the excision of WT 5’ fragment from the original clone. The list of primers used for cloning and site-directed mutagenesis is given in and the list of plasmids used in this study is given in Table S1.

### Cell culture and transfection

HEK293T and HeLa cells were cultured in DMEM high glucose medium (D7777, Sigma) supplemented with 10% FBS (Gibco) and antibiotic-antimycotic (Gibco), and maintained at 37°C under standard 5% CO2 conditions. Cells were transfected with the indicated plasmids using Lipofectamine 2000 (Thermo Fisher Scientific) or Lipofectamine 3000 (Thermo Fisher Scientific) as per the manufacturer’s instructions.

### Stable cell line generation

HeLa CRISPR-HIRA KO cells [23] were a kind gift from Dr. Geneviève Almouzni. These cells were transfected with H3.3-SNAP plasmid using Lipofectamine 3000 as per the manufacturer’s instructions. Stable transfectants expressing H3.3-SNAP were selected with Geneticin (700µg/ml). The expression of H3.3-SNAP was confirmed by western blot. For generating rescue cells that express WT, K600Q, and K600R HIRA, HeLa-HIRA KO cells were transfected with the corresponding plasmids, and the stable cells were selected with Geneticin (750µg/ml). The expression was confirmed using Western blot.

### H3.3-SNAP labeling *in vivo*

HeLa cells stably expressing H3.3-SNAP were platted on 24-well coverslips and transfected with WT, K600Q, and K600R mutants of HIRA using Lipofectamine 3000. Following (24 hours of transfection) the labeling of old/new histones was performed.

- **Labeling old histones:** To track old histones, the pulse-chase strategy was followed. The cells were incubated in a complete medium containing 2 μM SNAP-Cell TMR-Star (NEB) for 20 min to label all pre-existing H3.3-SNAP (pulse). After rinsing twice with PBS, the cells were incubated with a complete medium for 30 min to allow diffusion of excess SNAP-Cell TMR-Star. The cells were further incubated in the complete medium for 3 h or 9 h (chase).
- **Labeling new histones:** To track new histones, the quench-chase-pulse strategy was followed. The cells were incubated in a complete medium containing 10 μM SNAP-Cell Block (NEB) to block all available pre-existing H3.3-SNAP (quench), followed by two PBS washes and 30 min of incubation in complete medium to allow diffusion of the SNAP-Cell Block. The cells were further incubated in the complete medium for either 0 h (that is, background levels), or 2 h (chase), then performed TMR-Star labeling (pulse) as described above.
- **Extraction and fixation:** To visualize the chromatin-bound H3.3 and to remove the soluble pools of histones, Triton X-100 extraction was performed on the labeled cells before fixation. Pre-extraction was carried out by incubating the cells at room temperature for 5 min in 0.5% Triton in CSK buffer (10 mM PIPES (pH 7.0), 100 mM NaCl, 300 mM sucrose, 3 mM MgCl_2_), followed by two rapid rinses with CSK buffer and one PBS wash. Cells were then immediately fixed in 2% formaldehyde for 20 min at room temperature. After fixation, the cells were proceeded for HIRA immunofluorescence or stained with DAPI and mounted.

### Immunofluorescence

Post fixation, the cells were washed thrice with PBS and blocked for 1 hour in 5% BSA. Cells were incubated with HIRA antibody (Active motif) or other antibodies as mentioned at a concentration of 1:500 to 1:1000 overnight (12-14 hours). Cells were washed thrice with PBS and incubated with Alexa Fluor conjugated antibodies (Thermo-Fisher Scientific) for 2 hours at room temperature. Cells were washed three times with PBS and stained with DAPI (1:1000 of 1mg/ml). The coverslips were mounted in Vectashield on a glass slide.

### EU labeling

For nascent RNA labeling, cells were incubated with 10 μM EU (Click-iT™ RNA Alexa Fluor™ 488 Imaging Kit) for 30 min and the assay was performed as per the manufacturer’s instruction. Cells were then immediately fixed in 2% formaldehyde for 20 min at room temperature. After fixation, cells were washed in PBS, stained with DAPI, and mounted

### Microscopy and image analysis

Cells were imaged at 40X using an FV3000 laser scanning confocal microscope from Olympus Life Science. All experiments were repeated at least twice (N=2) and 150-250 cells per coverslips were analyzed for quantitation. The images were analyzed in FIJI (ImageJ) software to quantify fluorescence signal within the nuclei area.

### Cell culture Treatments

All treatments were given in high glucose medium supplemented with 10% FBS and antibiotic-antimycotic unless otherwise specified. For the heat shock paradigm, cells post-transfection were shifted to an incubator at 42°C for 90 minutes. After 90 minutes cells were shifted back to the incubator set at 37°C and allowed to recover for 4 hours. For the IFN paradigm, IFNγ was added to the cells at a concentration of 50ng/ml for 24 hours. Following the treatments, cells were collected in TriZol and proceeded for RNA isolation.

### RNA isolation, cDNA synthesis, and real-time PCR

Total RNA was isolated from cells using TriZol reagent. 1-2µg RNA was used to prepare cDNA using random hexamers and SuperScript-IV kit, as per the manufacturer’s protocol. Real time PCR (qPCR) was performed using KAPA SYBR® FAST Universal 2X qPCR Master Mix on Roche Light Cycler 480 II and LC 96 instruments. For normalization, levels of actin mRNA were used. The primers used for qPCR are given in Table S1.

### Cell fractionation

Cells were lysed in hypotonic lysis buffer (10 mM HEPES pH 7.5, 1.5 mM MgCl2, 10 mM KCl, 0.5 mM DTT, 1 mM PMSF, PhosSTOP, protease inhibitor cocktail) on ice for 15 minutes. Following the lysis, nuclei were pelleted by centrifugation at 2000 rpm for 15 minutes at 4**°**C. The supernatant was re-centrifuged at 12,000 rpm for 15 minutes at 4**°**C, the pellet (if any) was discarded and the supernatant obtained was collected as the cytoplasmic fraction. The nuclei were further rinsed with ice-cold hypotonic buffer twice and lysed in TNN lysis buffer (50mM Tris-HCl pH 7.5, 150mM NaCl, 0.9% NP 40, 1mM PMSF, PhosSTOP, protease inhibitor cocktail) for 15 minutes (incubation on ice). The lysate was further centrifuged at 12,000 rpm for 15 minutes at 4**°**C. The supernatant obtained was collected as the nucleoplasmic fraction. The pellet that contained the chromatin fraction was further rinsed with the TNN lysis buffer twice and lysed in RIPA buffer (50mM Tris pH 8.0, 150mM NaCl, 0.1% SDS, 0.5% Sodium deoxycholate, 1% Triton X 100, 0.1% SDS, 1 mM PMSF, Protease inhibitor cocktail), followed by sonication. After sonication the lysate was centrifuged at 12,000 rpm for 15 minutes at 4**°**C and the supernatant was collected as chromatin fraction. All the fractions were boiled in SDS gel loading buffer (50mM Tris-Cl pH 6.8, 2% SDS, 0.1% bromophenol blue, 10% glycerol, 100mM DTT) and loaded onto SDS-PAGE or used for immunoprecipitation with indicated antibodies. In the case of fractionation in high salt conditions, the same protocol was followed except that the TNN buffer contained 300 or 600mM NaCl (or indicated salt concentration) instead of 150mM.

### Immunoprecipitation

The TNN lysates were precleared with Protein A/G magnetic beads (Bio-Rad or Thermo Fisher Scientific) for 1 hour at 4**°**C on the rotor. After pre-clearing, the supernatant obtained was incubated with the indicated antibodies/antibody-conjugated beads overnight at 4**°**C on the rotor. The antibody-protein complexes were pulled down either directly (in cases where antibody-conjugated beads were used) or with protein-A/G beads (in cases where the antibody was incubated with the lysate overnight). The beads were washed thrice with the TNN lysis buffer for 10 minutes at 4**°**C on the rotor and boiled in SDS gel loading buffer for 15 minutes at 95**°**C.

### LC/MS-MS analysis of the acetylated proteome from mouse liver tissue

Mouse liver tissue was homogenized in lysis buffer (20 mM HEPES pH 8, 6 M Urea with protease inhibitors (Complete™, Roche) using a glass homogenizer, then sonicated with a Bioruptor (Diagenode). After clearing the lysates by centrifugation at 20000g for 15min, proteins were reduced with 5mM DTT and alkylated with 10mM iodoaceteamide. For enzymatic digestion, the urea concentration was brought to 1M by dilution of the samples in 50mM ammonium bicarbonate buffer, and proteins were digested with 1% (w/w) of trypsin overnight at 37°C. Tryptic peptides were desalted on C18 cartridges (Sep-Pak, Waters), and dried in a SpeedVac concentrator. Acetylated peptides were then enriched using agarose beads coupled with acetyl lysine antibody (ICP0388, ImmuneChem), by incubation in IAP Buffer (50 mM MOPS/NaOH pH 7.2, 10 mM Na2HPO4, 50 mM NaCl, Cell Signalling Technology), and eluted with 0.15% TFA. Tryptic peptides were dried and resuspended in 17 µl of 2% acetonitrile, 0.05% trifluoroacetic acid, and analyzed by nano-liquid chromatography (LC) coupled to tandem MS, using an UltiMate 3000 system (NCS-3500RS Nano/Cap System; Thermo Fisher Scientific) coupled to an Orbitrap Q Exactive Plus mass spectrometer (Thermo Fisher Scientific). Five microliters of each sample were loaded on a C18 precolumn (300 µm inner diameter × 5 mm, Thermo Fisher Scientific) in a solvent made of 2% acetonitrile and 0.05% trifluoroacetic acid, at a flow rate of 20 µl/min. After 5 min of desalting, the precolumn was switched online with the analytical C18 column (75 µm inner diameter × 50 cm, in-house packed with Reprosil C18) equilibrated in 95% solvent A (5% acetonitrile, 0.2% formic acid) and 5% solvent B (80% acetonitrile, 0.2% formic acid). Peptides were eluted using a 5%-50% gradient of solvent B over 120min at a flow rate of 300 nl/min. The mass spectrometer was operated in data-dependent acquisition mode with the Xcalibur software. MS survey scans were acquired with a resolution of 70,000 and an AGC target of 3e6. The 10 most intense ions were selected for fragmentation by high-energy collision induced dissociation, and the resulting fragments were analyzed at a resolution of 17500, using an AGC target of 1e5 and a maximum fill time of 100ms. Dynamic exclusion was used within 30 s to prevent repetitive selection of the same peptide. The raw MS data was processed with the Mascot (Matrix Science, version 2.8.0.1) search engine for database search against the Uniprot mouse reference proteome database plus a set of common contaminant proteins, using the following search parameters: Carbamidomethylation of cysteine was set as a fixed modification, whereas oxidation of methionine, protein N-terminal acetylation, and acetylation of lysine were set as variable modifications. Specificity of trypsin digestion was set for cleavage after K or R, and two missed trypsin cleavage sites were allowed. The mass tolerance was set to 10 ppm for precursors and 20mmu for fragments in MS/MS mode. Minimum peptide length was set to 7 amino acids. Mascot results were validated by the target decoy approach using the Proline software (Bouyssie et al, Bioinformatics, 2020) at both a peptide and protein false-discovery rate of 1%.

### Western Blotting

Equal amount of protein (50-100μg) or immunoprecipitation samples were run on SDS-PAGE and transferred to PVDF membrane (Millipore). The membranes were then blocked in 5% skimmed milk at room temperature for 1 hour on the rocker. Blots were cut according to molecular weights indicated by a pre-stained ladder (Abcam) and appropriate primary antibodies were added and incubated overnight (12-16 hours) at 4°C. Blots were washed and incubated with appropriate secondary antibodies. After the secondary antibody incubation, the blots were washed thrice and Supersignal West Pico PLUS or Supersignal West Femto ECL kit (Thermo Fisher Scientific) was used to detect the bands on GE Amersham Imager 600. All the antibodies, chemicals, and reagents used in this study are tabulated in Table S1.

### Quantification and statistical analysis

Data are expressed as means ± standard error of means (SEM). Statistical analyses were done using Graph Pad Prism (version 8.0). Student’s t-test and ANOVA were used to determine statistical significance. A value of p ≤ 0.05 was considered statistically significant. *p ≤ 0.05; **p ≤ 0.01; ***p ≤ 0.001.

## Acknowledgments

We thank Dr. Geneviève Almouzni, Dr. Dominique Ray-Gallet, and Audrey Forest for sharing HIRA CRISPR KO cells, critical inputs, and constructive discussions. We are grateful to Dr. Peter Adams and Dr. Ronen Marmorstein for sharing the UBN1 plasmids with us. We extend our acknowledgement to UK lab members for their critical inputs and discussions during the study.

## Funding

This research has been supported by the following funding sources: TIFR/DAE (19P0911), TIFR/DAE (19P0116), Department of Science and Technology JCB/2022/000036, Department of Biotechnology (BT/PR29878/PFN/20/1431/2018)

## Author contributions

Conceptualization: S.US., U.K-S., Methodology: S.US., M.L-P., A.G.P., U.K-S., Investigation: S.US., S.C., A.F., M.L-P., M.M., A.G.P., Visualization: S.US., S.C., A.F., M.L-P., M.M., A.G.P., Supervision: U.K-S., Writing—original draft: S.US., U.K-S.

## Competing interests

The authors declare that they have no competing interests.

## Data and materials availability

All data needed to evaluate the conclusions in the paper are present in the paper and/or the Supplementary Materials.

## Notes

### Competing Interest Statement

The authors have declared no competing interest.

## References

1. T. Kouzarides, Chromatin modifications and their function. Cell 128, 693–705 (2007).

2. G. Millán-Zambrano, A. Burton, A. J. Bannister, R. Schneider, Histone post-translational modifications - cause and consequence of genome function. Nat Rev Genet 23, 563–580 (2022).

3. S. Martire, L. A. Banaszynski, The roles of histone variants in fine-tuning chromatin organization and function. Nat Rev Mol Cell Biol 21, 522–541 (2020).

4. I. Maze, K. M. Noh, A. A. Soshnev, C. D. Allis, Every amino acid matters: essential contributions of histone variants to mammalian development and disease. Nat Rev Genet 15, 259–271 (2014).

5. G. D. Bowman, M. G. Poirier, Post-translational modifications of histones that influence nucleosome dynamics. Chem Rev 115, 2274–2295 (2015).

6. V. Sokolova, S. Sarkar, D. Tan, Histone variants and chromatin structure, update of advances. Comput Struct Biotechnol J 21, 299–311 (2023).

7. P. B. Talbert, S. Henikoff, Histone variants at a glance. J Cell Sci 134, (2021).

8. T. Yadav, J. P. Quivy, G. Almouzni, “Chromatin plasticity: A versatile landscape that underlies cell fate and identity” in *Science* (© 2018, American Association for the Advancement of Science., United States, 2018), vol. 361, pp. 1332–1336.

9. E. López-Jiménez, C. González-Aguilera, Role of Chromatin Replication in Transcriptional Plasticity, Cell Differentiation and Disease. Genes (Basel) 13, (2022).

10. S. B. Hake, C. D. Allis, Histone H3 variants and their potential role in indexing mammalian genomes: the “H3 barcode hypothesis”. Proc Natl Acad Sci U S A 103, 6428–6435 (2006).

11. D. Ray-Gallet, G. Almouzni, The Histone H3 Family and Its Deposition Pathways. Adv Exp Med Biol 1283, 17–42 (2021).

12. D. Filipescu, S. Müller, G. Almouzni, Histone H3 variants and their chaperones during development and disease: contributing to epigenetic control. Annu Rev Cell Dev Biol 30, 615–646 (2014).

13. K. Ahmad, S. Henikoff, The histone variant H3.3 marks active chromatin by replication-independent nucleosome assembly. Mol Cell 9, 1191–1200 (2002).

14. L. A. Banaszynski, C. D. Allis, P. W. Lewis, Histone variants in metazoan development. Dev Cell 19, 662–674 (2010).

15. L. Shi, H. Wen, X. Shi, The Histone Variant H3.3 in Transcriptional Regulation and Human Disease. J Mol Biol 429, 1934–1945 (2017).

16. E. Szenker, D. Ray-Gallet, G. Almouzni, The double face of the histone variant H3.3. Cell Res 21, 421–434 (2011).

17. P. Chen, J. Zhao, Y. Wang, M. Wang, H. Long, D. Liang, L. Huang, Z. Wen, W. Li, X. Li, H. Feng, H. Zhao, P. Zhu, M. Li, Q. F. Wang, G. Li, H3.3 actively marks enhancers and primes gene transcription via opening higher-ordered chromatin. Genes Dev 27, 2109–2124 (2013).

18. T. Tamura, M. Smith, T. Kanno, H. Dasenbrock, A. Nishiyama, K. Ozato, Inducible deposition of the histone variant H3.3 in interferon-stimulated genes. J Biol Chem 284, 12217–12225 (2009).

19. G. Banumathy, N. Somaiah, R. Zhang, Y. Tang, J. Hoffmann, M. Andrake, H. Ceulemans, D. Schultz, R. Marmorstein, P. D. Adams, Human UBN1 is an ortholog of yeast Hpc2p and has an essential role in the HIRA/ASF1a chromatin-remodeling pathway in senescent cells. Mol Cell Biol 29, 758–770 (2009).

20. S. J. Elsaesser, C. D. Allis, HIRA and Daxx constitute two independent histone H3.3-containing predeposition complexes. Cold Spring Harb Symp Quant Biol 75, 27–34 (2010).

21. T. S. Rai, A. Puri, T. McBryan, J. Hoffman, Y. Tang, N. A. Pchelintsev, J. van Tuyn, R. Marmorstein, D. C. Schultz, P. D. Adams, Human CABIN1 is a functional member of the human HIRA/UBN1/ASF1a histone H3.3 chaperone complex. Mol Cell Biol 31, 4107–4118 (2011).

22. C. Xiong, Z. Wen, J. Yu, J. Chen, C. P. Liu, X. Zhang, P. Chen, R. M. Xu, G. Li, UBN1/2 of HIRA complex is responsible for recognition and deposition of H3.3 at cis-regulatory elements of genes in mouse ES cells. BMC Biol 16, 110 (2018).

23. D. Ray-Gallet, M. D. Ricketts, Y. Sato, K. Gupta, E. Boyarchuk, T. Senda, R. Marmorstein, G. Almouzni, Functional activity of the H3.3 histone chaperone complex HIRA requires trimerization of the HIRA subunit. Nat Commun 9, 3103 (2018).

24. J. Torné, D. Ray-Gallet, E. Boyarchuk, M. Garnier, P. Le Baccon, A. Coulon, G. A. Orsi, G. Almouzni, Two HIRA-dependent pathways mediate H3.3 de novo deposition and recycling during transcription. Nat Struct Mol Biol 27, 1057–1068 (2020).

25. S. Ramazi, J. Zahiri, Posttranslational modifications in proteins: resources, tools and prediction methods. Database (Oxford) 2021, (2021).

26. T. Chattopadhyay, B. Maniyadath, H. P. Bagul, A. Chakraborty, N. Shukla, S. Budnar, A. Rajendran, A. Shukla, S. S. Kamat, U. Kolthur-Seetharam, Spatiotemporal gating of SIRT1 functions by O-GlcNAcylation is essential for liver metabolic switching and prevents hyperglycemia. Proc Natl Acad Sci U S A 117, 6890–6900 (2020).

27. T. Narita, B. T. Weinert, C. Choudhary, Functions and mechanisms of non-histone protein acetylation. Nat Rev Mol Cell Biol 20, 156–174 (2019).

28. C. Hall, D. M. Nelson, X. Ye, K. Baker, J. A. DeCaprio, S. Seeholzer, M. Lipinski, P. D. Adams, HIRA, the human homologue of yeast Hir1p and Hir2p, is a novel cyclin-cdk2 substrate whose expression blocks S-phase progression. Mol Cell Biol 21, 1854–1865 (2001).

29. X. Ye, B. Zerlanko, A. Kennedy, G. Banumathy, R. Zhang, P. D. Adams, Downregulation of Wnt signaling is a trigger for formation of facultative heterochromatin and onset of cell senescence in primary human cells. Mol Cell 27, 183–196 (2007).

30. J. H. Yang, T. Y. Song, C. Jo, J. Park, H. Y. Lee, I. Song, S. Hong, K. Y. Jung, J. Kim, J. W. Han, H. D. Youn, E. J. Cho, Differential regulation of the histone chaperone HIRA during muscle cell differentiation by a phosphorylation switch. Exp Mol Med 48, e252 (2016).

31. J. S. Lee, Z. Zhang, O-linked N-acetylglucosamine transferase (OGT) interacts with the histone chaperone HIRA complex and regulates nucleosome assembly and cellular senescence. Proc Natl Acad Sci U S A 113, E3213–3220 (2016).

32. Y. Chen, W. Zhao, J. S. Yang, Z. Cheng, H. Luo, Z. Lu, M. Tan, W. Gu, Y. Zhao, Quantitative acetylome analysis reveals the roles of SIRT1 in regulating diverse substrates and cellular pathways. Mol Cell Proteomics 11, 1048–1062 (2012).

33. C. B. Gocke, H. Yu, J. Kang, Systematic identification and analysis of mammalian small ubiquitin-like modifier substrates. J Biol Chem 280, 5004–5012 (2005).

34. B. T. Weinert, T. Narita, S. Satpathy, B. Srinivasan, B. K. Hansen, C. Schölz, W. B. Hamilton, B. E. Zucconi, W. W. Wang, W. R. Liu, J. M. Brickman, E. A. Kesicki, A. Lai, K. D. Bromberg, P. A. Cole, C. Choudhary, Time-Resolved Analysis Reveals Rapid Dynamics and Broad Scope of the CBP/p300 Acetylome. Cell 174, 231–244.e212 (2018).

35. L. Shi, B. P. Tu, Acetyl-CoA and the regulation of metabolism: mechanisms and consequences. Curr Opin Cell Biol 33, 125–131 (2015).

36. K. J. Menzies, H. Zhang, E. Katsyuba, J. Auwerx, “Protein acetylation in metabolism - metabolites and cofactors” in Nat Rev Endocrinol (England, 2016), vol. 12, pp. 43-60.

37. B. R. Zhou, H. Feng, R. Ghirlando, H. Kato, J. Gruschus, Y. Bai, Histone H4 K16Q mutation, an acetylation mimic, causes structural disorder of its N-terminal basic patch in the nucleosome. J Mol Biol 421, 30–37 (2012).

38. M. Li, J. Luo, C. L. Brooks, W. Gu, “Acetylation of p53 inhibits its ubiquitination by Mdm2” in J Biol Chem (United States, 2002), vol. 277, pp. 50607-50611.

39. X. Wang, J. J. Hayes, Acetylation mimics within individual core histone tail domains indicate distinct roles in regulating the stability of higher-order chromatin structure. Mol Cell Biol 28, 227–236 (2008).

40. K. Kamieniarz, R. Schneider, “Tools to tackle protein acetylation” in Chem Biol (United States, 2009), vol. 16, pp. 1027-1029.

41. H. Tagami, D. Ray-Gallet, G. Almouzni, Y. Nakatani, Histone H3.1 and H3.3 complexes mediate nucleosome assembly pathways dependent or independent of DNA synthesis. Cell 116, 51–61 (2004).

42. D. Ray-Gallet, J. P. Quivy, H. W. Silljé, E. A. Nigg, G. Almouzni, The histone chaperone Asf1 is dispensable for direct de novo histone deposition in Xenopus egg extracts. Chromosoma 116, 487–496 (2007).

43. C. M. English, N. K. Maluf, B. Tripet, M. E. Churchill, J. K. Tyler, ASF1 binds to a heterodimer of histones H3 and H4: a two-step mechanism for the assembly of the H3-H4 heterotetramer on DNA. Biochemistry 44, 13673–13682 (2005).

44. A. D. Goldberg, L. A. Banaszynski, K. M. Noh, P. W. Lewis, S. J. Elsaesser, S. Stadler, S. Dewell, M. Law, X. Guo, X. Li, D. Wen, A. Chapgier, R. C. DeKelver, J. C. Miller, Y. L. Lee, E. A. Boydston, M. C. Holmes, P. D. Gregory, J. M. Greally, S. Rafii, C. Yang, P. J. Scambler, D. Garrick, R. J. Gibbons, D. R. Higgs, I. M. Cristea, F. D. Urnov, D. Zheng, C. D. Allis, Distinct factors control histone variant H3.3 localization at specific genomic regions. Cell 140, 678–691 (2010).

45. D. Ray-Gallet, A. Woolfe, I. Vassias, C. Pellentz, N. Lacoste, A. Puri, D. C. Schultz, N. A. Pchelintsev, P. D. Adams, L. E. Jansen, G. Almouzni, Dynamics of histone H3 deposition in vivo reveal a nucleosome gap-filling mechanism for H3.3 to maintain chromatin integrity. Mol Cell 44, 928–941 (2011).

46. J. Torné, G. A. Orsi, D. Ray-Gallet, G. Almouzni, Imaging Newly Synthesized and Old Histone Variant Dynamics Dependent on Chaperones Using the SNAP-Tag System. Methods Mol Biol 1832, 207–221 (2018).

47. C. Sadamitsu, T. Nagano, Y. Fukumaki, A. Iwaki, Heat shock factor 2 is involved in the upregulation of alphaB-crystallin by high extracellular potassium. J Biochem 129, 813–820 (2001).

48. Q. Jiang, Z. Zhang, Y. Hu, Y. Ma, Function of Hsf1 in SV40 T antigen transformed HEK293T cells. Mol Med Rep 10, 3139–3144 (2014).

49. W. Siwek, S. S. H. Tehrani, J. F. Mata, L. E. T. Jansen, Activation of Clustered IFNγ Target Genes Drives Cohesin-Controlled Transcriptional Memory. Mol Cell 80, 396–409.e396 (2020).

50. L. J. Benson, Y. Gu, T. Yakovleva, K. Tong, C. Barrows, C. L. Strack, R. G. Cook, C. A. Mizzen, A. T. Annunziato, Modifications of H3 and H4 during chromatin replication, nucleosome assembly, and histone exchange. J Biol Chem 281, 9287–9296 (2006).

51. A. Loyola, T. Bonaldi, D. Roche, A. Imhof, G. Almouzni, PTMs on H3 variants before chromatin assembly potentiate their final epigenetic state. Mol Cell 24, 309–316 (2006).

52. P. Cramer, Organization and regulation of gene transcription. Nature 573, 45–54 (2019).

53. E. Vojnic, B. Simon, B. D. Strahl, M. Sattler, P. Cramer, Structure and carboxyl-terminal domain (CTD) binding of the Set2 SRI domain that couples histone H3 Lys36 methylation to transcription. J Biol Chem 281, 13–15 (2006).

54. K. O. Kizer, H. P. Phatnani, Y. Shibata, H. Hall, A. L. Greenleaf, B. D. Strahl, A novel domain in Set2 mediates RNA polymerase II interaction and couples histone H3 K36 methylation with transcript elongation. Mol Cell Biol 25, 3305–3316 (2005).

55. D. K. Pokholok, C. T. Harbison, S. Levine, M. Cole, N. M. Hannett, T. I. Lee, G. W. Bell, K. Walker, P. A. Rolfe, E. Herbolsheimer, J. Zeitlinger, F. Lewitter, D. K. Gifford, R. A. Young, Genome-wide map of nucleosome acetylation and methylation in yeast. Cell 122, 517–527 (2005).

56. S. Henikoff, M. M. Smith, Histone variants and epigenetics. Cold Spring Harb Perspect Biol 7, a019364 (2015).

57. Z. A. Gurard-Levin, J. P. Quivy, G. Almouzni, Histone chaperones: assisting histone traffic and nucleosome dynamics. Annu Rev Biochem 83, 487–517 (2014).

58. M. Tyagi, N. Imam, K. Verma, A. K. Patel, Chromatin remodelers: We are the drivers!! Nucleus 7, 388–404 (2016).

59. S. J. Elsässer, S. D’Arcy, Towards a mechanism for histone chaperones. Biochim Biophys Acta 1819, 211–221 (2013).

60. F. Mattiroli, S. D’Arcy, K. Luger, The right place at the right time: chaperoning core histone variants. EMBO Rep 16, 1454–1466 (2015).

61. M. D. Ricketts, R. Marmorstein, A Molecular Prospective for HIRA Complex Assembly and H3.3-Specific Histone Chaperone Function. J Mol Biol 429, 1924–1933 (2017).

62. C. Das, J. K. Tyler, M. E. Churchill, The histone shuffle: histone chaperones in an energetic dance. Trends Biochem Sci 35, 476–489 (2010).

63. L. De Koning, A. Corpet, J. E. Haber, G. Almouzni, Histone chaperones: an escort network regulating histone traffic. Nat Struct Mol Biol 14, 997–1007 (2007).

64. C. M. Hammond, C. B. Strømme, H. Huang, D. J. Patel, A. Groth, Histone chaperone networks shaping chromatin function. Nat Rev Mol Cell Biol 18, 141–158 (2017).

65. P. Grover, J. S. Asa, E. I. Campos, H3-H4 Histone Chaperone Pathways. Annu Rev Genet 52, 109–130 (2018).

66. S. K. Pradhan, T. Su, L. Yen, K. Jacquet, C. Huang, J. Côté, S. K. Kurdistani, M. F. Carey, EP400 Deposits H3.3 into Promoters and Enhancers during Gene Activation. Mol Cell 61, 27–38 (2016).

67. A. García Del Arco, B. A. Edgar, S. Erhardt, In Vivo Analysis of Centromeric Proteins Reveals a Stem Cell-Specific Asymmetry and an Essential Role in Differentiated, Non-proliferating Cells. Cell Rep 22, 1982–1993 (2018).

68. E. H. Zion, D. Ringwalt, K. Rinaldi, E. W. Kahney, Y. Li, X. Chen, Old and newly synthesized histones are asymmetrically distributed in Drosophila intestinal stem cell divisions. EMBO Rep 24, e56404 (2023).

69. A. Wenger, A. Biran, N. Alcaraz, A. Redó-Riveiro, A. C. Sell, R. Krautz, V. Flury, N. Reverón-Gómez, V. Solis-Mezarino, M. Völker-Albert, A. Imhof, R. Andersson, J. M. Brickman, A. Groth, Symmetric inheritance of parental histones governs epigenome maintenance and embryonic stem cell identity. Nat Genet 55, 1567–1578 (2023).

70. Y. M. Xu, J. Y. Du, A. T. Lau, Posttranslational modifications of human histone H3: an update. Proteomics 14, 2047–2060 (2014).

71. M. E. Gerritsen, A. J. Williams, A. S. Neish, S. Moore, Y. Shi, T. Collins, CREB-binding protein/p300 are transcriptional coactivators of p65. Proc Natl Acad Sci U S A 94, 2927–2932 (1997).

72. F. Wang, C. B. Marshall, M. Ikura, Transcriptional/epigenetic regulator CBP/p300 in tumorigenesis: structural and functional versatility in target recognition. Cell Mol Life Sci 70, 3989–4008 (2013).

73. T. Zhang, W. L. Kraus, SIRT1-dependent regulation of chromatin and transcription: linking NAD(+) metabolism and signaling to the control of cellular functions. Biochim Biophys Acta 1804, 1666–1675 (2010).

74. Y. Dai, D. V. Faller, Transcription Regulation by Class III Histone Deacetylases (HDACs)-Sirtuins. Transl Oncogenomics 3, 53–65 (2008).

75. B. Nashun, P. W. Hill, S. A. Smallwood, G. Dharmalingam, R. Amouroux, S. J. Clark, V. Sharma, E. Ndjetehe, P. Pelczar, R. J. Festenstein, G. Kelsey, P. Hajkova, Continuous Histone Replacement by Hira Is Essential for Normal Transcriptional Regulation and De Novo DNA Methylation during Mouse Oogenesis. Mol Cell 60, 611–625 (2015).

76. Q. Kong, L. A. Banaszynski, F. Geng, X. Zhang, J. Zhang, H. Zhang, C. L. O’Neill, P. Yan, Z. Liu, K. Shido, G. D. Palermo, C. D. Allis, S. Rafii, Z. Rosenwaks, D. Wen, Histone variant H3.3-mediated chromatin remodeling is essential for paternal genome activation in mouse preimplantation embryos. J Biol Chem 293, 3829–3838 (2018).

