## Supplementary figures and images for "Biased recruitment of H3.3 by HIRA is dictated by de-/acetylation and determines transcription memory and response"

### Supplementary Figure 1

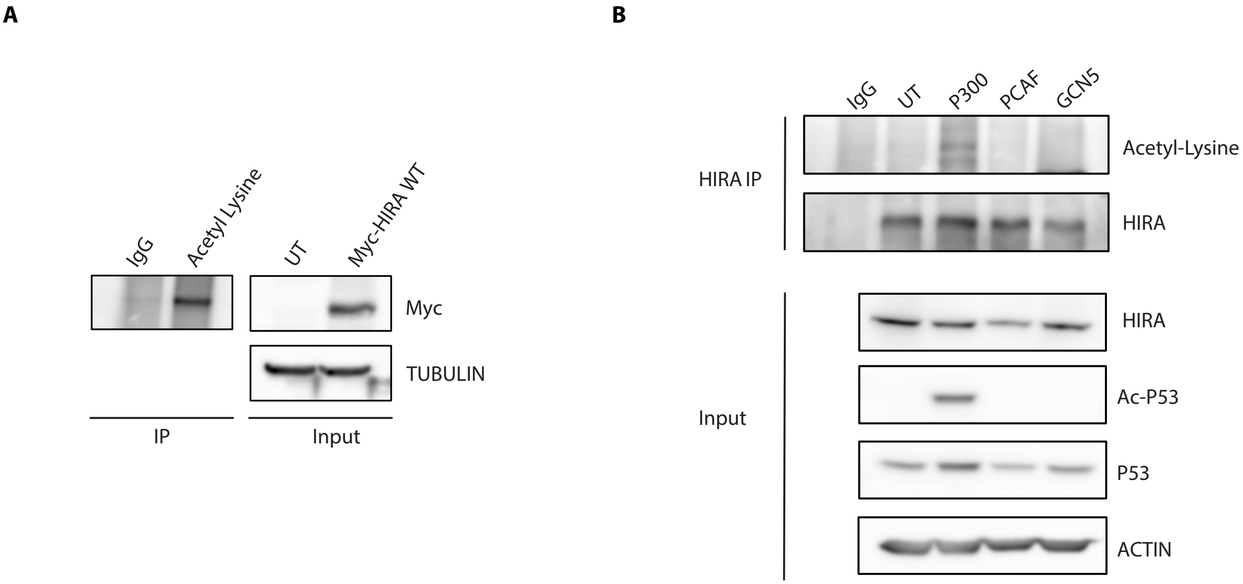


Supplementary Figure-S1

### Supplementary Figure 2

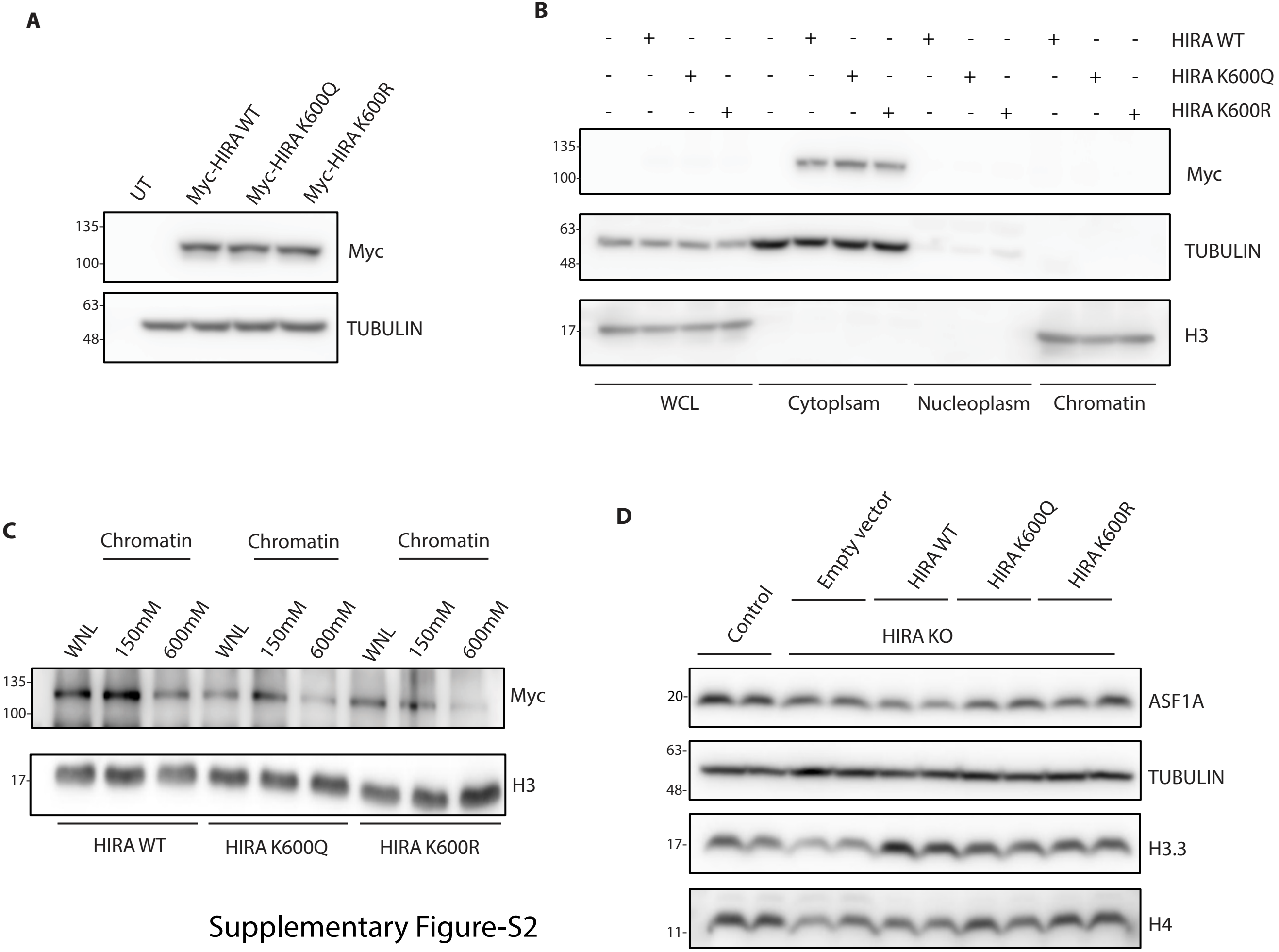


Supplementary Figure-S2

### Supplementary Figure 3

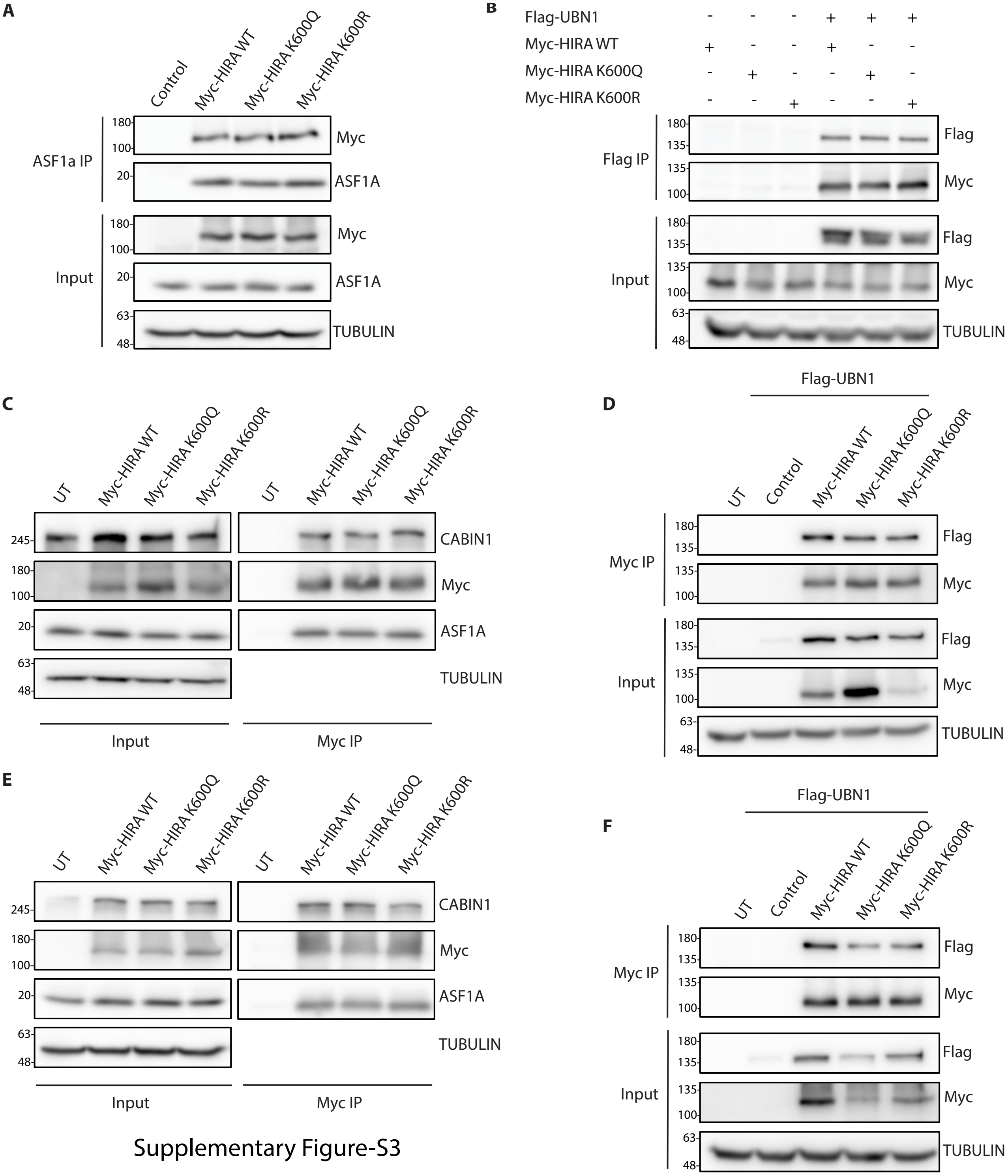


Supplementary Figure-S3

### Supplementary Figure 4

#### Supplementary Figure-S4

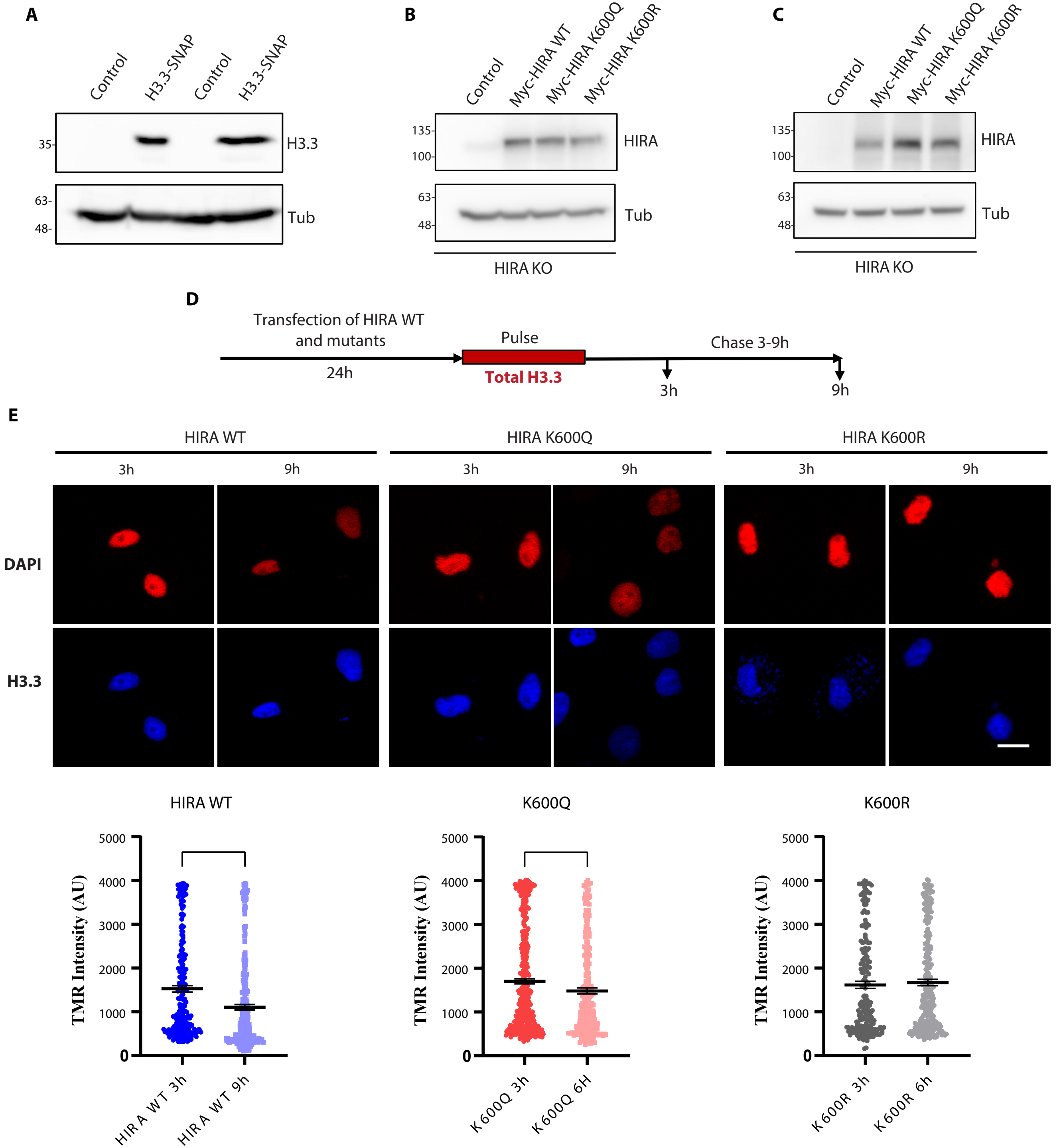

### Supplementary Figure 5

Supplementary Figure-S5

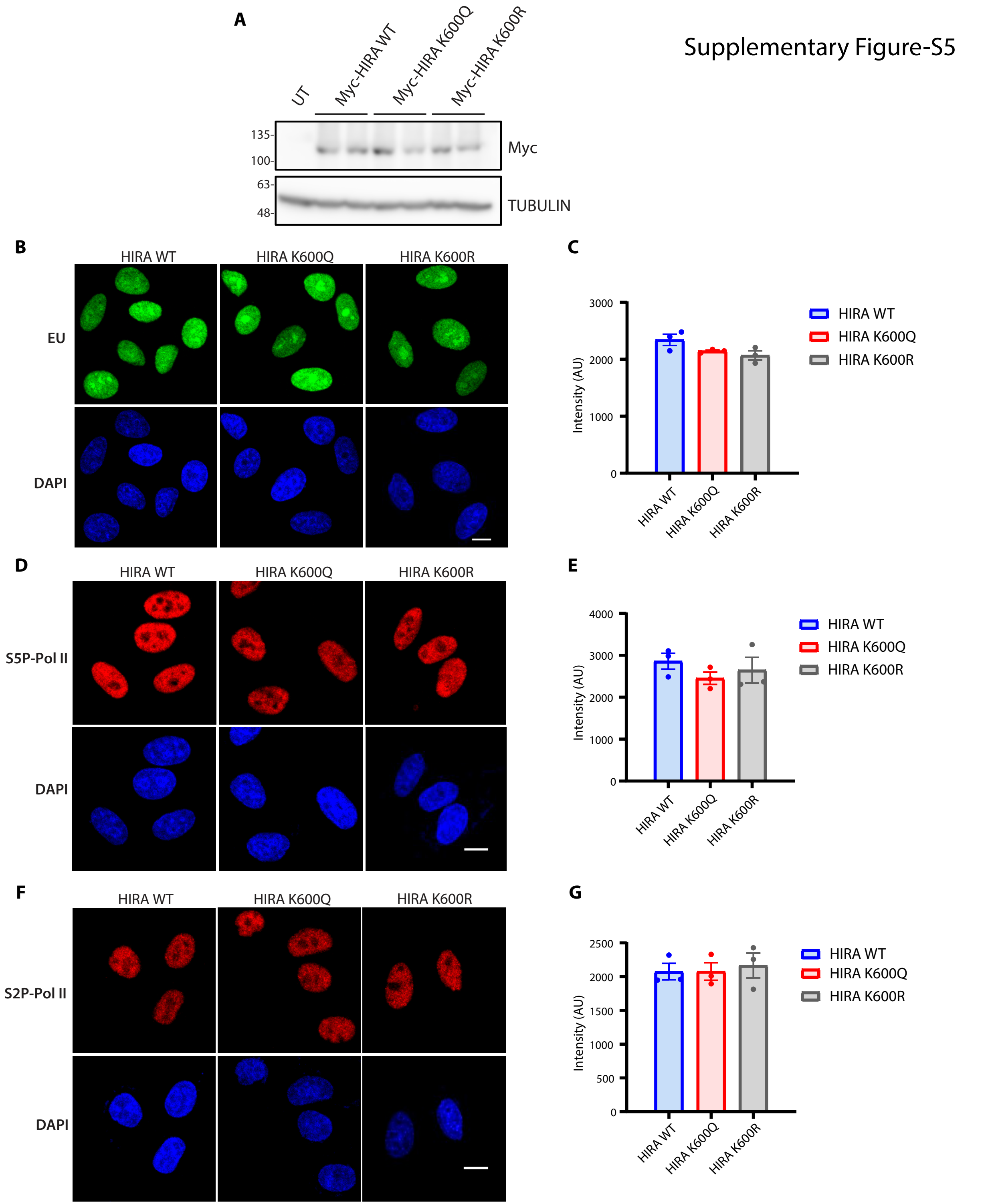
