## Supplementary Information for "Biased recruitment of H3.3 by HIRA is dictated by de-/acetylation and determines transcription memory and response"

**Fig. S1. Mammalian HIRA is acetylated (A)** HEK293T cells transfected with Myc-HIRA WT followed by immunoprecipitation (IP) assay performed using pan-acetyl lysine antibody and western blot using Myc antibody to detect acetylated HIRA (N=2). **(B)** HEK293T cells transfected with P300, PCAF, or GCN5 followed by IP assay using HIRA antibody and western blot analysis using pan-acetyl lysine antibody (N=2).

**Fig. S2. Acetylation alters HIRA chromatin association (A)** HEK293T cells transfected with Myc-HIRA WT, K600Q, and K600R followed by western blot assay using Myc antibody to compare expression of HIRA constructs (N=3). **(B)** HEK293T cells transfected with Myc-HIRA WT, K600Q, and K600R followed by biochemical fractionation and western blot analysis to assay for Myc-HIRA levels in whole cell lysate (WCL), cytoplasm, nucleoplasm, and chromatin fractions (N=3). **(C)** HEK293T cells transfected with Myc-HIRA WT, K600Q, and K600R followed by biochemical fractionation with nuclear lysis done at low (150mM) and high salt (600mM) conditions. Western blot analysis of chromatin fraction to assay for Myc-HIRA levels (N=3). **(D)** Representative blots for levels of ASF1A and H3.3 in control, HIRA KO, and HIRA KO cells rescued with HIRA WT, K600Q, and K600R (n=2 N=3).

**Fig. S3. HIRA complexation** **is independent of acetylation (A)** HEK293T cells transfected with Myc-HIRA WT, K600Q, and K600R followed by coIP assay using ASF1A antibody and western blot for Myc HIRA levels (N=2). **(B)** HEK293T cells co-transfected with Flag-UBN1 and Myc-HIRA WT, K600Q, and K600R followed by coIP assay using Flag beads and western blot for Myc-HIRA levels to assay for interaction (N=2). **(C)** HEK293T cells transfected with Myc-HIRA WT, K600Q, and K600R followed by coIP assay in TNN buffer with 300mM salt using Myc beads and western blot to assay for interactions with ASF1A and CABIN1 (N=2). **(D)** HEK293T cells co-transfected with Flag-UBN1 and Myc-HIRA WT, K600Q, and K600R followed by coIP assay in TNN buffer with 300mM salt using Myc beads and western blot for Flag to assay for interaction with Flag-UBN1 (N=2). **(E)** HEK293T cells transfected with Myc-HIRA WT, K600Q, and K600R followed by coIP assay in TNN buffer with 600mM salt using Myc beads and western blot to assay for interactions with ASF1A and CABIN1 (N=2). **(F)** HEK293T cells co-transfected with Flag-UBN1 and Myc-HIRA WT, K600Q, and K600R followed by coIP assay in TNN buffer with 600mM salt using Myc beads and western blot for Flag to assay for interaction with Flag-UBN1 (N=2).

**Fig. S4. HIRA K600R exhibits enhanced H3.3 recycling activity (A)** Representative blots showing H3.3-SNAP levels in HIRA-KO cells stably expressing H3.3-SNAP. **(B)** Western blot showing HIRA WT, K600Q, and K600R levels in *de novo* H3.3 deposition paradigm. **(C)** Western blot showing HIRA WT, K600Q, and K600R levels in recycling paradigm. **(D)** Schematic depicting the experimental paradigm for assessing H3.3 recycling activity. **(E)** Representative immunofluorescence images showing H3.3-SNAP (red) and DAPI (blue) in Hela cells stably expressing SNAP-H3.3. Cells were transfected with Myc-HIRA WT, K600Q, and K600R followed by *in vivo* pulse labeling of pre-existing H3.3-SNAP (at 0h) and a chase period of 3h and 9h. The quantifications are represented as mean ± S.E.M with all the data points shown (N=2, n>150). Statistical significance was calculated using a t test (*, p < 0.05; **, p < 0.01; ***, p < 0.001). Scale bars -10 µm.

**Fig. S5. HIRA rescue cells exhibit no change in global transcription and levels of phosphorylated RNA Pol-II (A)** Western blot showing Myc-HIRA WT, K600Q, and K600R levels in heat shock paradigm. **(B-C)** Representative images showing EU labeling (green) that marks nascent RNA in HIRA KO cells rescued with HIRA WT, K600Q, and K600R using **(B)** immunofluorescence assay and **(C)** its quantification. **(D-G)** Representative images for Ser 5 phosphorylated Pol II, **(D)** S5P-Pol II and **(F)** S2P-Pol II in HIRA KO cells rescued with HIRA WT, K600Q, and K600R using immunofluorescence assay and the quantification respectively **(E and G)**. Data represented as mean ± S.E.M. (N=2, n=3). Statistical significance was calculated using one-way ANOVA with Tukey’s test for multiple comparisons between groups (*, p < 0.05; **, p < 0.01; ***, p < 0.001). Scale bars -10 µm.

| **Primers used for cloning and site-directed mutagenesis** | | | |
| --- | --- | --- | --- |
| **Primer name** | **Sequence (5’ to 3’)** | | |
| hH3.3 SNAP FP | AATGGATCCGGCGCGCCATGGCTCGTACAAAGCAGACTGCCC | | |
| hH3.3 SNAP RP | ATAAAGCTTGAATTCAGCACGTTCTCCACGTATGCGGCGTG | | |
| hHIRA Common FP | AAAGAATTCATGAAGCTCCTGAAGCCGACC | | |
| hHIRA K600Q RP | AAAGTCGACCTAGGGCCTCAGCTCTTGCACAAG | | |
| hHIRA K600R RP | AAAGTCGACCTAGGGCCTCAGCTCTCTCACAAG | | |
| hHIRA WT FL RP | GCTGTAGGACTCCCTGTTCATCCAGCTGAAA | | |
| **Primers used for qPCR analysis** | | | |
| hHSP70 qFP | GTGTGTAACCCCATCATCAG | | |
| hHSP70 qRP | CACAGGAAATTGAGAACTGAC | | |
| hHSP27 qFP | AGTCCAACGAGATCACCATC | | |
| hHSP27 qRP | GTGGTTGCTTTGAACTTTATTTG | | |
| hGBP1 qFP | GTGGAACGTGTGAAAGCTGA | | |
| hGBP1 qRP | CAACTGGACCCTGTCGTTCT | | |
| hGBP5-qFP | TTCAATTTGCCCCGTCTGTG | | |
| hGBP5 qRP | AGGCAGTGTTTCAAGTTGGG | | |
| hHLA-DRA qFP | GAAAGCAGTCATCTTCAGCGTT | | |
| hHLA-DRA qRP | GAGGCATTGGCATGGTGATAAT | | |
| hCD74 qFP | TGGGAGGTGACTGTCAGTTTG | | |
| hCD74 qRP | AGGCTTTTCCATCCTGGTGAC | | |
| **Plasmids used in the study** | | | |
| **Plasmid name** | **Catalogue number** | **Gifted by** | |
| pAd-Track Flag-SIRT1 | 8438 (Addgene) | Pere Puigserver | |
| pAdEasy Flag GCN5 | 14106 (Addgene) | Pere Puigserver | |
| pCMVb P300 | 10717 (Addgene) | William Sellers | |
| pCI flag PCAF | 8941 (Addgene) | Yoshihiro Nakatani | |
| pLU Flag UBN1 |  | Dr. Peter Adams and Dr. Ronen Marmorstein | |
| pSNAP_f_ | N9183S (NEB) |  | |
| pSNAP_f_ -H3.3 |  | Cloned (in this study) | |
| pCMV-Tag3B Myc HIRA WT |  | Cloned (in this study) | |
| pCMV-Tag3B Myc HIRA K600Q |  | Cloned (in this study) | |
| pCMV-Tag3B Myc HIRA K600R |  | Cloned (in this study) | |
| **Antibodies used in the study** | | | |
| **Antibody name** | **Company** | **Catalogue number** | **Dilution** |
| Mouse IgG-HRP | Sigma | A9044 | 1:8000 |
| Rabbit IgG-HRP | Sigma | A0545 | 1:8000 |
| Alexa Fluor 488-conjugated anti-mouse antibody | Thermo Fisher Scientific | A28175 | 1:500 |
| Actin | Sigma | A1978 | 1:20000 |
| Tubulin | Sigma | T8328 | 1:8000 |
| FLAG M2 | Sigma | F1804 | 1:1000 |
| Myc | Cell Signaling Technology | 2276 | 1:1000 |
| P53 | Cell Signaling Technology | 2524 | 1:1000 |
| Acetyl p53 | Cell Signaling Technology | 2570 | 1:1000 |
| UBN1 | Abcam | ab101282 | 1:1000 |
| CABIN1 | Cell Signaling Technology | 12660 | 1:1000 |
| ASF1A | Cell Signaling Technology | 2990 | 1:1000 |
| H3.3 | Abcam | ab176840 | 1:2000 |
| H3 | Abcam | ab1791 | 1:8000 |
| H4 | Abcam | ab31830 | 1:1000 |
| H3K36me3 | Diagenode | CS-058-100 | 1:2000 |
| H3K4me3 | Diagenode | C15410003 | 1:2000 |
| H3K9Ac | Diagenode | C15410004 | 1:2000 |
| Acetyl-lysine | Cell Signaling Technology | 9814 | 1:1000 |
| HIRA | Active motif | 39557 | 1:500 |
| HA-Tag | Abcam | ab9110 | 1:1000 |
| Lamin B1 | Abcam | ab16048 | 1:1000 |
| **Reagents/chemicals used in the study** | | | |
| **Chemical/Reagent name** | | **Company** | **Catalog number** |
| RNaseOUT™ Recombinant Ribonuclease Inhibitor | | Invitrogen | 10777019 |
| Random Hexamer (50uM) | | Invitrogen | N8080127 |
| TriZol | | Invitrogen | 15596018 |
| DEPC treated water | | Invitrogen | AM9915G |
| UltraPure Distilled Water | | Invitrogen | 10977015 |
| dNTP Mix | | Genei | 652400021730 |
| DAPI | | Roche | 10236276001 |
| High Glucose DMEM | | Sigma-Aldrich | D7777 |
| Fetal Bovine Serum | | Gibco™ | 10270106 |
| Antibiotic Antimycotic Solution | | Sigma-Aldrich | A5955 |
| EDTA | | Sigma-Aldrich | E9884 |
| Bovine serum albumin | | MP Biomedicals | 160069 |
| DTT | | Sigma-Aldrich | 578517 |
| Tris Base | | Himedia | MB029 |
| HEPES | | USB | 16926 |
| 20% SDS | | Himedia | ML007 |
| Magnetic Protein A beads | | Biorad | #1614013 |
| Lipofectamine™ 3000 Transfection Reagent | | Invitrogen | L3000001 |
| Lipofectamine™ 2000 Transfection Reagent | | Invitrogen | 11668019 |
| IGEPAL® CA-630 | | Sigma-Aldrich | I3021 |
| Triton X-100 | | Sigma-Aldrich | T8787 |
| Sodium deoxycholate | | Sigma-Aldrich | SRE0046 |
| Sodium Bicarbonate | | Himedia | TC230 |
| Sodium Chloride | | MP™ | 194848 |
| cOmplete™, EDTA-free Protease Inhibitor Cocktail | | Roche | COEDTAF-RO |
| PhosSTOP™ | | Roche | PHOSS-RO |
| Phenylmethanesylfonyl fluoride (PMSF) | | Sigma-Aldrich | P7626 |
| Bicinchoninic Acid solution | | Sigma-Aldrich | B9643 |
| Tween 20 | | Himedia | TC287 |
| PVDF Membrane | | Millipore | IPVH00010 |
| Skim milk powder | | Himedia | GRM1254 |
| Paraformaldehyde | | Sigma-Aldrich | 158127 |
| Chemiluminescence detection kit | | Thermo Scietific | 34095, 34080 |
| SuperScript-IV RT kit | | Invitrogen | 18090200 |
| KAPA SYBR® FAST Universal 2X qPCR Master Mix | | Roche | SFUKB |
| SNAP-Cell^®^ Block | | New England Biolabs | S9106S |
| SNAP-Cell^®^ TMR-Star | | New England Biolabs | S9105S |
| Click-iT™ RNA Alexa Fluor™ 488 Imaging Kit | | Invitrogen | C10329 |

**Table S1 Key Resource Table**
